# Core-N-glycans are atypically abundant at the neuronal surface and regulate glutamate receptor signaling

**DOI:** 10.1101/2024.03.25.586577

**Authors:** Chun-Lei Zhang, Cédric Moutoussamy, Matthieu Tuffery, Alexandre Varangot, Rebecca Piskorowski, Cyril Hanus

## Abstract

Neurotransmitter receptors, like most surface proteins, are extensively modified by covalent addition of N-glycans during their synthesis. Surprisingly, the most abundant N-glycans in the mammalian brain are core-glycans, sugars that typically earmark immature intracellular proteins in non-neuronal cells. The function of these glycans in neurons is yet largely unknown. To address this, we combined conditional gene knockout, mass spectrometry, quantitative imaging and electrophysiological recordings in cultured neurons and brain slices. We show that core-glycans are expressed at high levels at the neuronal surface, indicating expression on functional proteins. Focusing on excitatory synapses, we found that core-glycans reduce dendritic spine density and synaptic AMPA receptor expression but are overall sufficient to sustain functional synapses. Our results indicate that core-glycans slow the desensitization of AMPA receptor complexes and reduce NMDA receptor signaling at synapses. Core-glycans hence impair NMDA receptor-dependent synaptic plasticity, unraveling a previously unrecognized role for N-glycosylation in regulating synaptic composition and transmission efficacy.

## Introduction

N-glycans are found on more than 75% of membrane proteins and regulate protein assembly, stability and physical interactions with their ligands and partner proteins (1). Protein glycotypes are in some instance extremely diverse (1), potentially affecting protein chemical composition and function to a large degree. Akin to epigenetic mechanisms, N-glycosylation is modulated by cell environment and metabolism (2). N-glycan alterations are reported in an increasing number of brain pathologies, notably Alzheimer’s disease and schizophrenia (3–5), and are emerging as a promising and still largely untapped source of biomarkers.

Our previous work and more recent studies have shown that hundreds of neuronal membrane proteins are core-glycosylated (6–9) - *i.e.*, display N-glycans that have not been fully processed in the Golgi apparatus - contrasting with other tissues where mature surface proteins typically do not display such glycans (1, 6). Blocking N-glycan maturation in neural stem cells favors differentiation into neurons (10, 11), thus indicating that core-glycosylation plays an important role in the acquisition of neuronal identity.

Core-glycans are present on numerous neurotransmitter receptors and notably glutamate AMPA receptors, that mediate most of the fast excitatory synaptic transmission in the mammalian brain. As shown by deglycosylation and molecular-shift assays, these receptors coexist at the neuronal surface as standard and core-glycosylated glycoforms (6). Consistent with N-glycans role in controlling nascent proteins interactions with ER chaperon proteins (1), mutagenesis and heterologous expression have shown that specific N-glycosylation sites of AMPA receptor subunits are required for proper protein folding, assembly and surface expression (12). However, it is not yet known whether the acquisition of specific glycosylation profiles confer specific functional properties to AMPA receptor complexes.

Here, we used pharmacological and genetic approaches to prevent the maturation of core-glycans into “mature” N-glycans in the neuronal Golgi to address their function. By combining mass spectrometry, quantitative imaging and electrophysiological recordings in primary hippocampal neuron cultures and hippocampal slices, we show that core-glycans are among the most abundant N-glycans of the neuronal surface and regulate AMPA receptor desensitization and expression at synapses. Surprisingly, while core-glycans are sufficient for proper function of surface AMPA receptors at the plasma membrane, they attenuate long-term synaptic potentiation through impairment of NMDA receptor signaling. Altogether, these results provide a novel experimental and conceptual framework to understand how N-glycan processing regulates ionotropic glutamate receptors and synaptic plasticity.

## Results

### Core-glycans are atypically prevalent at the neuronal surface

Several in-depth MS studies of brain glycopeptides have been published over the last decade (4, 7–9, 13), providing a wealth of information on the exact composition and abundance of brain N-glycans. We reanalyzed these datasets to facilitate direct cross comparisons (see Methods). Consistent with our previous work with lectins (6) these data show that core-glycans are not only present but are the most abundant N-glycans in human and mouse brains (Figure 1B and Supplemental Figures S1A-S1C). Similar high abundance was observed in purified mouse synapses (see “synaptosomes” in Figure 1C and Supplemental Figures S1D), indicating that core-glycans are found on functional (synaptic) proteins. Because synaptosomes also contain endomembranes (14), it cannot be ruled out that these core-glycans may specifically originate from intracellular proteins. To address this, we purified surface membrane proteins after surface biotinylation of cultured rat neurons and analyzed their glycan composition by liquid-chromatography matrix assisted laser desorption/ionization time-of-flight (LC-MALDI-TOF) MS (Figure 1D and Supplemental Figure S2). Although their relative abundance slightly differed from total mouse brain and synaptosomes, we found that core-glycans were indeed expressed at high levels at the neuronal surface (Figure 1D).

**Figure 1.**
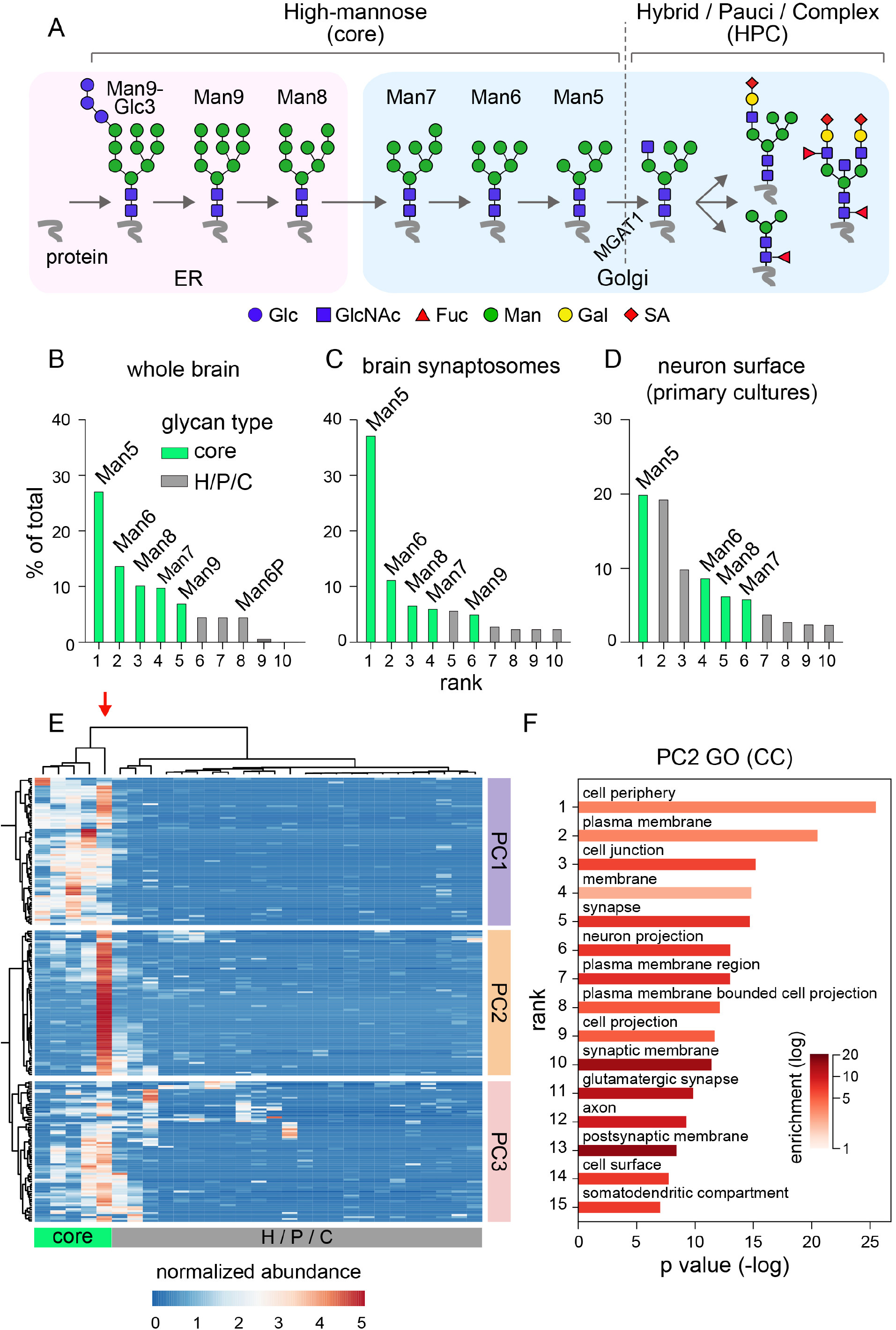
Prevalence of core-N-glycans in the brain and at the neuronal surface, see also Supplementary Figures S1-S5. **A**) Simplified scheme of N-glycan processing along the secretory pathway. Note the variations of the number of mannose residues (green disks) after exit from the endoplasmic reticulum (ER) and before processing by a-1,3-mannosyl-glycoprotein 2-b-N-acetylglucosaminyltransferase 1 (MGAT1 and subsequent maturation into hybrid/pauci/complex (HPC) N-glycans in the Golgi apparatus. Glc, glucose; GlcNAc, N-acetyl-glucosamine; Fuc, fucose; Man, mannose; Gal, galactose; SA, sialic acid. **B**-**D**) Occurrence of core-glycans (green) and HPC N-glycans (grey) among the top ten most abundant N-glycans identified by mass spectrometry in total mouse brain (B), mouse synaptosomes (C) and purified rat neuronal surface proteins (D). In the three types of samples, note the prevalence of Man5 and to a lesser extent Man6, Man8 and Man7. Plots in B and C were generated from raw data in ref. [(7)] and [(13)], respectively. **E)** Cluster map of mouse brain N-glycans (columns) plotted against peptide abundance (rows). Clustering was performed using Euclidian distance and Ward’s method. Generated from raw data in [(7)]. Note the segregation of core *versus* HPC N-glycans at the first bifurcation level (see dendrogram at the top of the graph) and the segregation of glycoproteins into 3 main clusters (PC1-PC3, see dendrogram on the left side of the graph). Man5 is shown among N-glycans by a red arrow. **F)** Gene ontology (GO) analysis showing the most enriched GO terms among proteins of cluster 2 shown in E compared to whole mouse proteome (see Methods). Shown are enrichment p-value color coded for fold enrichment of the first 15 “cellular components” GO terms (GO:CC). Note the occurrence of terms related to synaptic transmission and glutamatergic synapses.

Interestingly, GlcNAc-2-Mannose-5 (Man5) and to a lesser extent Man6, Man8 and Man7 – *i.e.* glycan types typically found on proteins after ER-exit but before maturation into hybrid, pauci or complex N-glycans (HPC) in the GA (Figure 1A) (15) – were among the most abundant N-glycans and were found in virtually the same ranking order in total brain, synaptosomes and surface samples (Figure 1B-1D, Supplemental Figure S1). Using clustering methods, we found that core-glycan expression was the main determinant of the segmentation of brain glycoproteins in 2 main distinct glycotypes (core-glycans *vs* HPC N-glycans) (Figure 1E). These glycotypes separated glycoproteins in 3 main families, with, at the first bifurcation level, proteins expressing relatively high levels of all core-glycans (protein cluster 1); then at the second bifurcation level two protein groups defined by a high expression of Man5 alone (protein cluster 2) or more diverse N-glycan types and a higher expression of some HPC glycans (protein cluster 3) (Figures 1E). As shown by gene ontology (GO) analysis of protein cluster 2, we found that Man5 expression is particularly high for postsynaptic proteins (Figure 1F, see also Figure 1C).

As shown by analysis of liver glycopeptides, high-levels of core-glycans in total cellular membranes are not specific to the brain (Supplemental Figure S3A). However, we found that Man5, Man6 and Man7 – the 3 N-glycans generated from Man8 in the GA before maturation into HPC sugars according to textbooks (15) - specifically stood out in the brain (Supplemental Figures S3A and S3B), indicating a tissue-specific regulation of this group of core-glycans.

Thus, core-glycans - most notably Man5 - are atypically prevalent on surface neuronal glycoproteins and in particular postsynaptic proteins.

### Neuron-specific knockout of MGAT1 enables experimental control of neuronal N-glycosylation

To experimentally increase Man5 abundance in neurons for functional studies, we used a conditional-knockout (cKO) mouse model to prevent neuronal expression of a-1,3-mannosyl-glycoprotein 2-b-N-acetylglucosaminyltransferase 1 (MGAT1) (16), the enzyme that controls the first step of core-glycan maturation into HPC N-glycans in the Golgi apparatus (Figure 1A) (15). Neuron-specific MGAT1 KO was performed by transduction of homozygous MGAT1^flx^ mouse brains or cultured neurons with adeno-associated virus (AAVs) enabling expression of fluorescent cell fills and Cre recombinase (Cre) under a neuron specific promoter (synapsin1, Supplemental Figure S4A). Resulting changes of N-glycosylation were assessed in cultured MGAT1^flx^ neurons transduced with AAVs encoding GFP or GFP plus Cre by far-western blotting with lectins (Supplemental Figure S4B) and electrospray-ionisation (LC-ESI) MS (Supplemental Figures S4C-SE and S5). As expected, MGAT1 knockout increased the expression of core-glycans, most prominently Man5, and decreased the expression of hybrid/complex N-glycans. Of note, the rank and relative expression levels of surface core-glycans in mouse neurons differed from rat (Figure 1D), likely reflecting differences in MS assays (LC-MALDI-TOF *vs* LC-ESI-MS-MS, see Discussion) and species.

Thus, MGAT1 cKO increases neuronal core-glycans - mostly in the form of Man5 - at the expense of mature N-glycans, enabling experimental control of neuronal N-glycosylation.

### Core-glycans regulate dendritic spine density and AMPA receptor expression at synapses

Previous studies with glycosydase inhibitors in cultured neurons showed that core-glycosylated proteins are functional and are sufficient to sustain synapse maintenance (6). To confirm this *ex vivo*, we assessed dendritic spine morphology in CA1 hippocampal pyramidal neurons, a well-established proxy to monitor changes of synaptic composition and activity (17, 18). To this aim, AAVs encoding GFP alone or GFP plus Cre under a synapsin1 gene promoter were stereotaxically injected in the dorsal hippocampus of 7 to 9 weeks-old MGAT1^flx^ mice (see Methods. Figure 2A and Supplemental Figure S6). Spine imaging and 3D reconstructions were performed in CA1 pyramidal neuron secondary and tertiary dendrites in *stratum radiatum* in fixed brain samples prepared 8 to 9 weeks after AAV transduction (Figures 2B, 2C). We found that spine density was ∼25% lower in Cre-expressing neurons (Figures 2C, 2D and Supplemental Figures S7 and S8) but was still within the range of reported physiological values (19). MGAT1 KO did not alter the proportion of main spine morphological types (Figures 2E), suggesting that reduction of spine density was not caused by impairment of spine maturation. Spine heads were, however, significantly enlarged by MGAT1 cKO (Figure 2F), perhaps indicating compensatory mechanisms counterbalancing the reduction of spine density. Although the density of GFP expressing neurons varied from one animal to another, Cre expressing neurons showed no sign of degeneration such as pyknotic bodies or severed dendrites (not shown).

**Figure 2.**
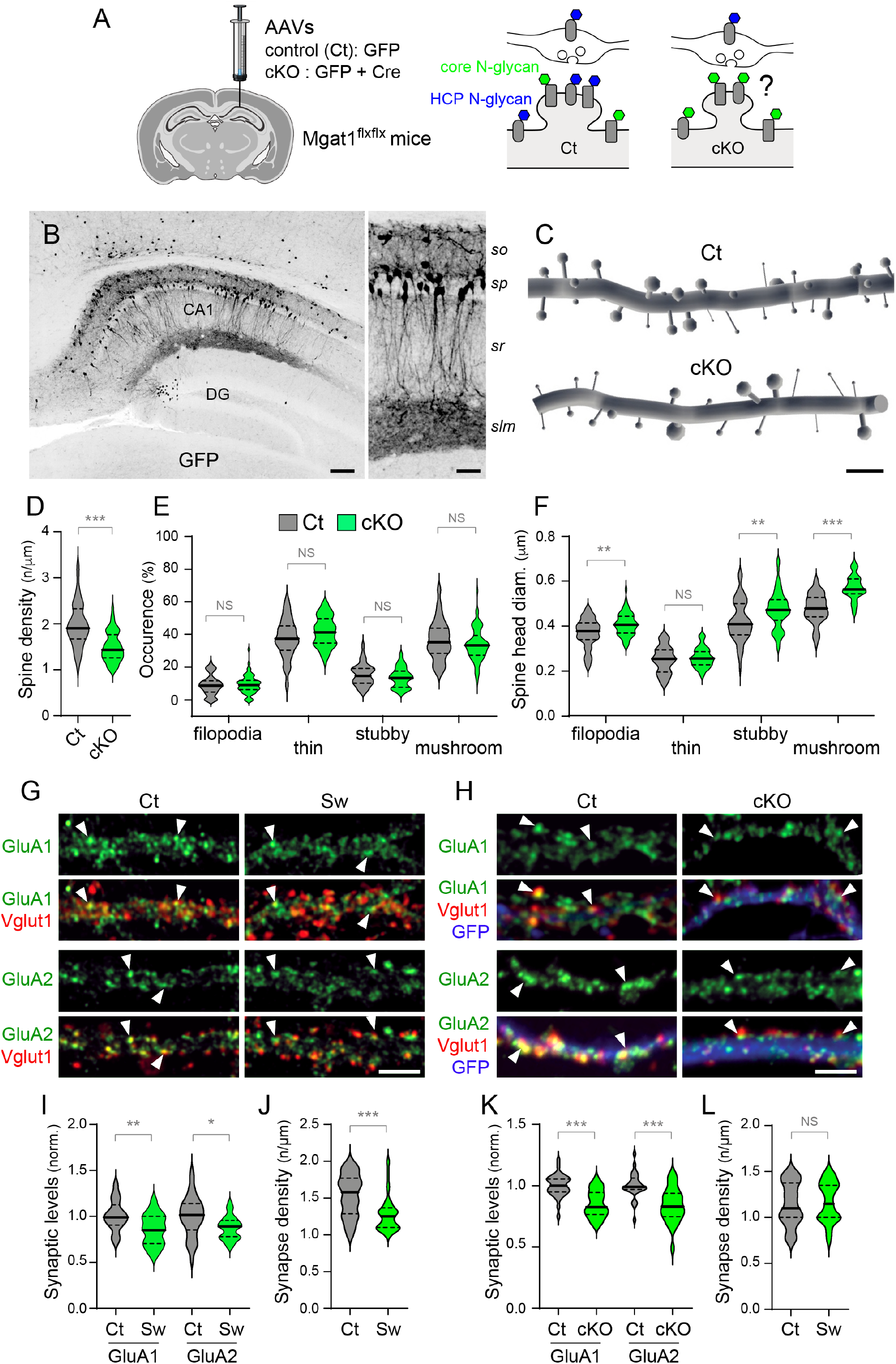
Core-glycans regulate dendritic spine density and AMPA receptor expression at synapses, see also Supplementary Figures S6-S9. **A**-**F**) Impact of Mgat1 conditional knockout (cKO) on spine morphology in CA1 pyramidal neurons *in vivo*. **A**) Mgat1 cKO was achieved by stereotaxic injection of AAVs encoding GFP±Cre under a synapsin 1 promoter in the dorsal hippocampus of 7-8 weeks-old Mgat1^flx/flx^ mice. The impact of preventing the biogenesis of HPC N-glycans on spine morphology was then assessed 8 weeks post AAV injection in fixed brain sections. **B**) Example confocal micrograph of GFP expressing neurons at low and higher magnification. Scale bars 100 or 50 μm. DG, dentate gyrus; *so*, *stratum oriens*, *sp*, *stratum pyramidale*; *sr*, *stratum radiatum*; *slm*, *stratum lacunosum moleculare*. **C**) Examples of spine reconstructions in secondary and tertiary dendrites of Mgat^flx^ in CA1 pyramidal neurons expressing GFP (control, Ct) or GFP+Cre (cKO). Scale bar 4 μm. **D**-**F**) Average spine density (D), spine type distribution (E) and spine head volumes (F) in control and Mgat1 KO neurons showing increased spine density and spine head volume in Mgat1 KO neurons. Shown in D-E are violin plots with median and interquartile range displayed as thick and dotted lines, respectively (n=46-60 dendrites, 5 mice per group). **, p<0.01; ***, p<10^-4^; NS, not significant; Kruskal-Wallis and Dunn’s *post-hoc* tests. **G**-**L**) Impact of Swainsonine (Sw) and Mgat1 KO on AMPA receptor expression at synapses. **G**,**H**) Confocal fluorescent micrographs of cultured hippocampal rat (G) and Mgat1^flx^ mouse neurons (H) after immunolabeling of AMPA GluA2 (green) and VGlut1 (red) in control conditions or after inhibition of core-glycan maturation with Sw or Mgat1 cKO. GFP fluorescence in mouse neurons is shown in blue. Arrows indicate synaptic receptor puncta. Scale bar 5 μm. **I**-**L**) Integrated fluorescence of GluA1 and GluA2 synaptic puncta (H, K) and density of “dendritic” VGlu1 puncta (J,L) in control neurons (Ct) and in cells chronically treated with Sw (Sw, I,J) or transduced with Cre-encoding AAVs (cKO, K-L). Shown are violin plots with median and interquartile range displayed as thick and dotted lines, respectively (n=28-65 cells, 3 experiments). *, p<0.05; **, p<0.01; ***, p<10^-4^; NS, not significant; Kruskal-Wallis and Dunn’s *post-hoc* tests. In G-L, note the reduction of AMPA receptor expression at synapses and the contrasted effect of Sw and Mgat1 KO on Vglut1 puncta density.

Changes of spine morphology are directly linked to synaptic composition and, in particular, to the number of synaptic AMPA receptors (20). We thus assessed whether increasing core-glycosylation modified AMPA receptor expression. To this end, we performed immuno-labeling experiments to compare the synaptic quantities of GluA1 and GluA2 (the two main subunits of AMPA receptors in the hippocampus)(21) in cultured mouse MGAT1 KO neurons and rat neurons chronically treated with swainsonine (Sw), an inhibitor preventing the maturation of core-glycans in the GA (Supplemental Figure S9A and Figures 2G-2L). We first verified that both manipulations modified AMPA receptor glycosylation in consistent ways using molecular shift assays after deglycosylation. As expected, chronic Sw treatment and MGAT1 KO both inhibited the maturation of core-glycosylated receptors into mature glycotypes (Supplemental Figure S9B). Receptor expression was then assessed by quantitative imaging at synapses visualized by accumulation of the presynaptic vesicular glutamate transporter 1 (VGLUT1). We found that Sw and MGAT1 KO both induced a ∼20% reduction of AMPA receptor expression at synapses (Figures 2G, 2H, 2I, 2K). Sw induced a mild albeit significant reduction of VGLUT1 puncta density (Figure 2J), that, in contrast, was not detected after MGAT1 cKO (Figure 2L).

In parallel, we performed surface biotinylation experiments and found that increasing core-glycosylation did not impair AMPA receptor expression at the plasma membrane (Supplementary Figures S9C and S9D). We thus conclude that although mature N-glycans are required for normal synaptic expression of AMPA receptors, core-glycans are sufficient to sustain the bulk of AMPA expression at the neuronal surface and synapses.

### Core-glycans regulate the electrophysiological properties of AMPA receptor complexes

Mannose-binding lectins have long been used to experimentally block the desensitization of AMPA and kainate receptors (22, 23). This suggests that polypeptide domains that control the desensitization of these receptors are core-glycosylated and that this core-glycosylation regulate receptor desensitization. As a first step towards addressing this, we measured AMPA miniature excitatory synaptic currents (mEPSCs) in cultured rat neurons after chronic treatment with Sw (Figures 3A and 3B). Consistent with immunolabeling experiments (Figures 2G, 2J), Sw induced a ∼20% decrease in mEPSC amplitude (Figure 3B), confirming reduced synaptic expression of AMPA receptors. Sw induced a 3-fold slowing of current decay yet had no effect on rise time or frequency (Figure 3B), thus indicating that core-glycosylation regulates receptor desensitization. To rule out that this effect on current decay was due to the N-glycosylation status of AMPA receptor complexes and not to indirect regulations by unrelated proteins, we recorded glutamate-induced currents in outside-out patches (Figures 3C and 3D). In this case, prior exposure to Sw did not significantly alter the amplitude of evoked currents (Figure 3D). In contrast, pre-treatment with Sw markedly slowed down current decay (Figure 3D), consistent with results obtained for synaptic currents.

**Figure 3.**
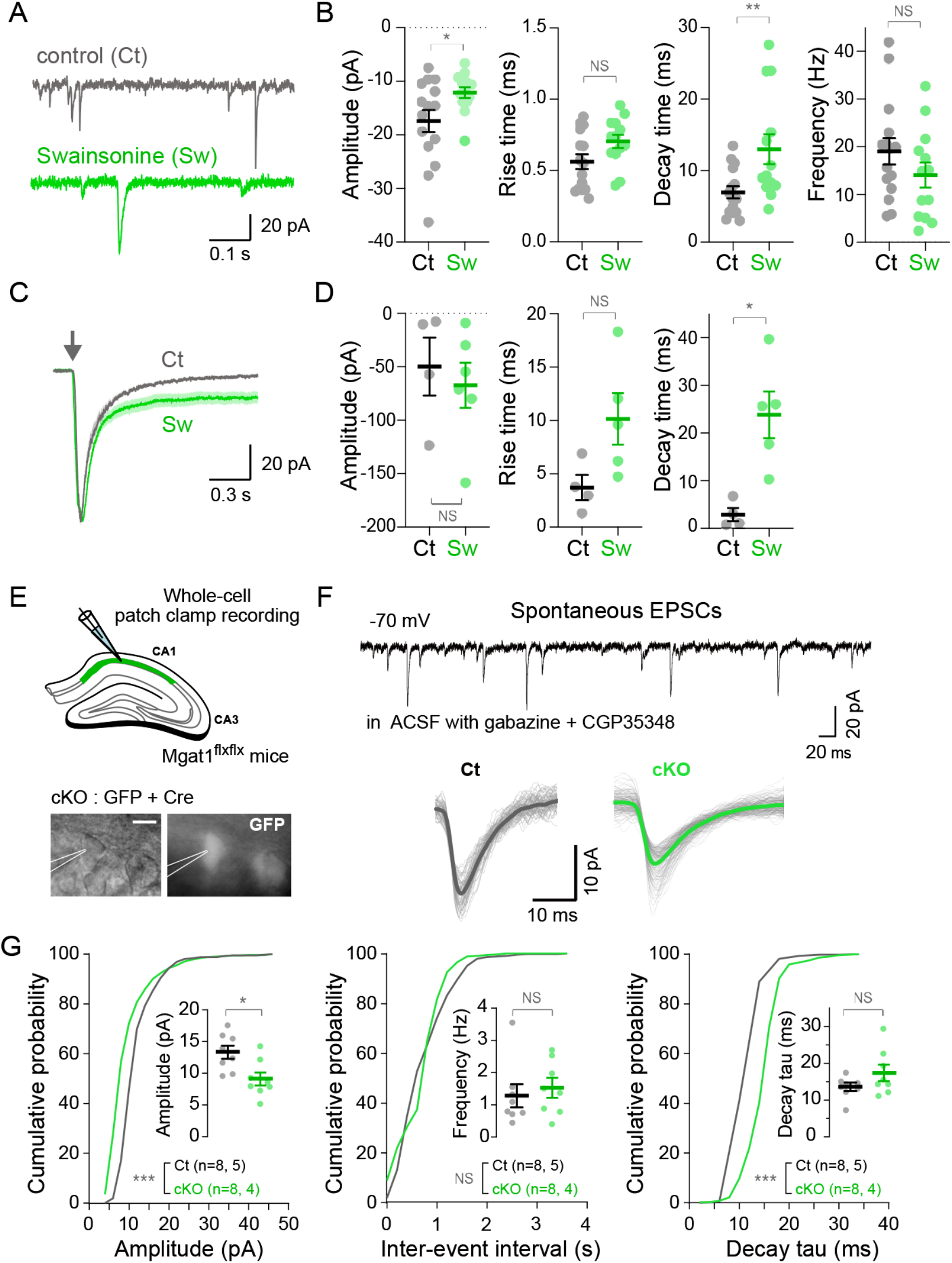
Core-glycans regulate AMPA receptor signaling. **A,B**) Patch-clamp recording of miniature AMPA excitatory postsynaptic currents (EPSCs) in rat neurons cultured with Swainsonine (Sw). **A**) Example traces from neurons in control condition (Ct) or after treatment with Sw (Sw). **B**) Average EPSC peak amplitude, decay tau, rise time and frequency in “Ct” and “Sw” neurons (means ± SEM, n=13-15 in 3 independent experiments). *, p<0.05; **, p <0.01; NS, non-significant; *Man-Whitney test*. **C**,**D**) Evoked AMPA receptor currents in outside-out membrane patches. **C**) Example traces from “Ct” or “Sw” neurons. Stimulation with glutamate is indicated by an arrow. **D**) Average current peak amplitude and rise and decay tau. Means ± SEM, n=4-6 in 3 independent experiments. *, p<0.05; NS, non-significant; *Man-Whitney test*. **E**-**G**) Patch-clamp recording of spontaneous EPSCs in acute slices prepared after *in vivo* injections of GFP (Ct) or GFP+Cre (cKO) AAVs. **E**) Scheme of experimental setup and microgaphs of a recorded neuron expressing GFP. **F**) Example traces recorded in “Ct” or “cKO” neurons. **G**) Distribution of current peak amplitude, decay tau and inter-event interval (M) in “Ct” and “cKO” groups. ***, p<10^-4^; NS, not significant; Kolmogorov-Smirnov’s test. Average values (means ± SEM) are shown in insets (n=8 recordings from 4-5 animals per group). ***, p <0.001; *, p<0.05; NS, non-significant; Man-Whitney tests. In A-G, note that Sw and Mgat1 cKO decrease EPSC amplitude and markedly slow down current decay.

To address whether core-glycans also affected synaptic currents in intact brain circuits, we recorded spontaneous AMPA-receptor mediated EPSCs (sEPSCs) in hippocampal CA1 pyramidal neurons in acute slices prepared from homozygous MGAT1^flx^ mouse brains 3 to 6 weeks after unilateral stereotaxic injection of AAVs encoding either GFP alone or GFP plus Cre in the dorsal hippocampus (Figures 3E-3G). Consistent with results in cultured neurons, neuron specific KO of MGAT1 resulted in a significant decreased of sEPSC amplitude and reduced decay kinetics but had no effect on sEPSC frequency (Figure 3G).

Thus, core-glycans slow the desensitization of AMPA receptor responses, hence indicating that the molecular complexes formed by specific AMPA receptor glycotypes have specific electrophysiological properties and signaling properties.

### Core-glycans reduce synaptic NMDA receptor signaling and impair long-term synaptic potentiation

Next, we addressed whether core-glycans had a different impact on synaptic NMDA and AMPA receptor responses and monitored evoked synaptic currents in acute hippocampal slices. The dorsal hippocampus of homozygous MGAT1^flx^ mice was unilaterally injected with AAVs encoding GFP alone or GFP plus Cre. Synaptic transmission was evoked by stimulating Schaeffer collateral inputs in CA1 *stratum radiatum.* Whole-cell patch clamp recordings were performed in visually identified CA1 pyramidal neurons that were either uninfected, expressing GFP alone, or expressing GFP and Cre (Figure 4A. see Methods for details). Recordings evoked synaptic transmission of of MGAT1 cKO neurons revealed a lower NMDA/AMPA ratio of evoked responses as compared to controls (Figure 4B), while the paired-pulse ratio was not altered (Figure 4C). Importantly, MGAT1 KO did not change either the decay timecourse nor the rise time of evoked NMDA receptor component of EPSCs (Supplemental Figures S10A-S10C). The input-output relationship of evoked EPSCs revealed that AMPA receptor response was not altered (Figure 4D), in contrast to NMDA receptor currents, which were strongly reduced in MGAT1 cKO neurons (Figure 4E).

**Figure 4:**
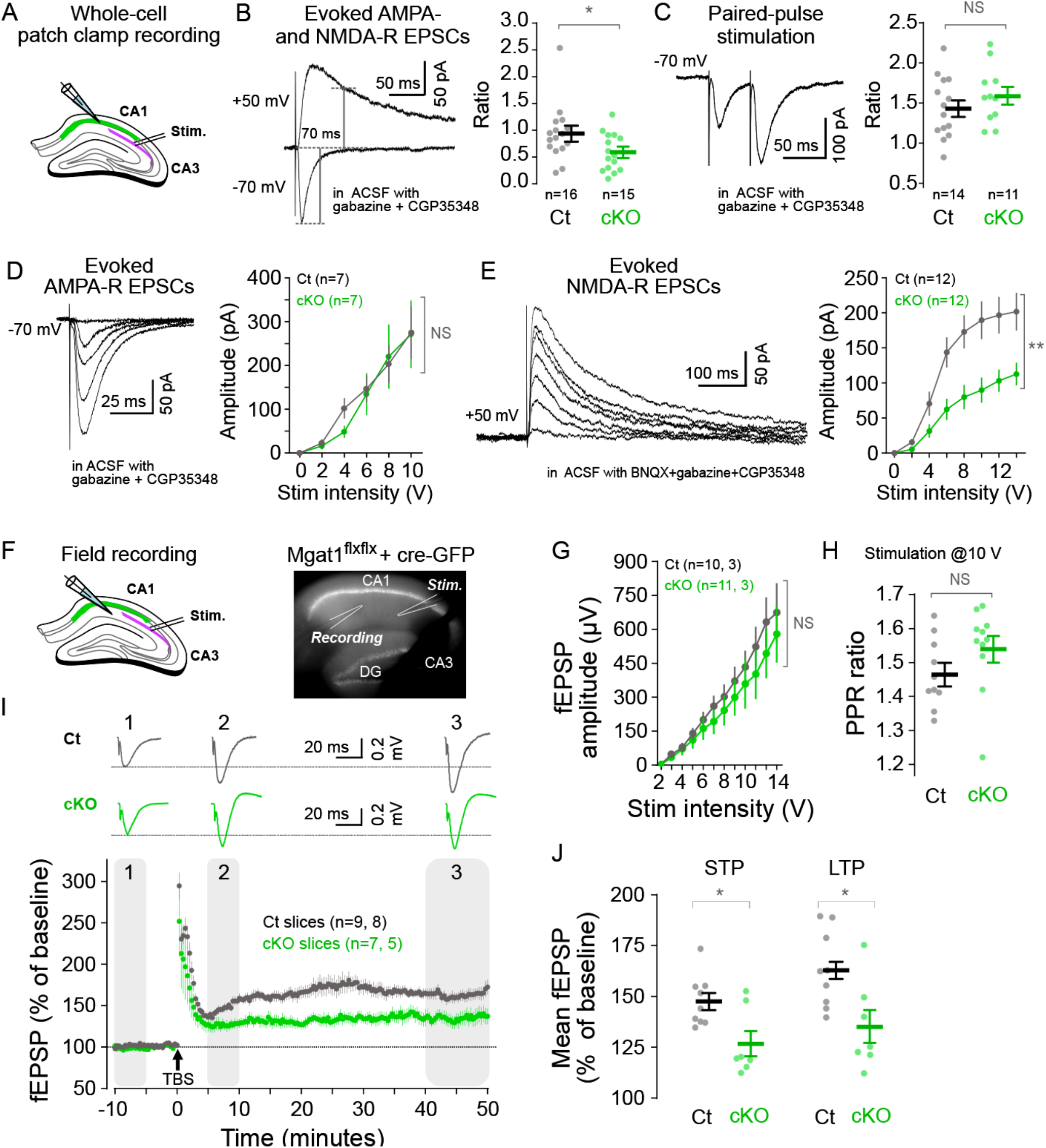
Core-glycans attenuate long-term synaptic potentiation and reduce NMDA receptor expression at the neuronal surface, see also Supplementary Figures S10. **A-E)** Patch clamp analysis of evoked excitatory postsynaptic currents (EPSCs) in control (Ct) and Mgat1 KO (cKO) neurons in acute slices. **A)** Scheme of experimental setup showing position of stimulating and recording electrodes. **B**) Example traces of evoked AMPA and NMDA receptor EPSCs and average NMDA to AMPA ratio in “Ct” and “cKO” neurons showing Mgat1 cKO impact on NMDA and AMPA receptor contribution to EPSCs. N=15-16 recordings from 5-7 animals per group. **C**) Example traces and average paired pulse ratio showing that Mgat1 cKO does not alter paired-pulse facilitation. N=11- 14 recordings from 3-6 animals per group. **D**,**E**) Example traces and average AMPA (D, N=7 recordings from 3-4 animals per group) and NMDA receptor EPSC amplitude (E, N=12 recordings from 5-6 animals per group) at varying stimulation intensities. Note that Mgat1 cKO selectively decreases evoked NMDA receptor currents in basal conditions. Average values in B-E are shown as means ± SEM. **, p <0.01; *, p<0.05; NS, non-significant; *Man-Whitney test (B, C) and two-way ANOVA (D, E)*. **F**-**J**) Long term potentiation of field excitatory postsynaptic potentials (fEPSPs) in control (Ct) and Mgat1 KO (cKO) CA1 pyramidal neurons in acute brain slices. **F)** Scheme and example fluorescence micrograph of experimental setup showing position of recorded GFP expressing neurons and position of stimulating and recording electrodes. **G**,**H**) Plot of fEPSP amplitude as a function of stimulation intensity (G) and paired-pulse ratio (200 ms interval) (H) in “Ct” and “cKO” slices showing that the two parameters are unaffected by Mgat1 cKO. N=10-11 slices from 3 animals per group. **I**,**J**) Example traces (I) and average of EPSP slopes (means ± SEM) in control and Mgat1 KO neurons (J) 5 to 10 min (short-term potentiation or STP) and 40 to 50 min after stimulation (long term potentiation) by tetanus bursts (TBS in D) showing the attenuation of LTP in Mgat1 KO neurons. N=7-9 slices from 5-8 animals per group. Shown in G-J are means±SEM. *, p<0.05; NS, non-significant; Man-Whitney’s test (H, J) and two-way ANOVA (G).

We then determined whether this decrease of synaptic NMDA receptor signaling affected synaptic plasticity. To this aim, we measured the long-term synaptic potentiation (LTP) at CA3-CA1 synapses in acute hippocampal slices following induction by theta burst stimulation (TBS, Figures 4F-4J), a widely studied form of postsynaptic plasticity induced by NMDA receptor activation (24). As for whole-cell patch clamp recording, the dorsal hippocampus of homozygous MGAT1^flx^ mice was unilaterally-injected with AAVs encoding GFP alone or GFP plus Cre. Synaptic transmission was evoked by stimulating Schaeffer collateral inputs in CA1 *stratum radiatum* and field excitatory post synaptic potentials (fEPSPs) recorded in regions of CA1 that were either uninfected, expressing GFP alone, or expressing GFP and Cre (Figure 4F). We found no difference in the LTP magnitude between untransfected slices or slices expressing GFP alone and pooled this data. When compared to slices expressing Cre, we found that both the early and late phases of LTP were significantly reduced (Figures 4I and 4J). Attenuated LTP in MGAT1 cKO slices occurred without concomitant variation in fiber volley amplitude nor paired-pulse ratio over the course of the experiments (Supplemental Figures S10D-E). Consistent with whole cell recordings (Figures 4C and 4D), MGAT1 cKO did not significantly affect the input-output relationship and paired-pulse ratio of fEPSP measured before LTP induction (Figures 4G and 4H).

These results thus indicate that MGAT1 cKO reduces NMDA synaptic responses and hence impairs synaptic plasticity.

## Discussion

Accumulating evidence shows that core-glycans, most notably Man5, are atypically prevalent in the mammalian brain (4, 7–9, 13). We show here that these glycans are present on *surface expressed* neuronal proteins and impact synaptic transmission by regulating the function of two key synaptic neurotransmitter receptors.

Changes of the N-glycosylation of synaptic proteins, including glutamate receptors, have been reported in multiple debilitating brain diseases, in particular schizophrenia and Alzheimer’s disease (AD)(3–5). Interestingly, recent studies have reported concomitant increases of the levels of MGAT1 mRNA and the expression of hybrid/complex N-glycans in the brain AD patients (5, 25). It is not clear at present whether these changes participate to the pathogenesis of the disease or merely reflect compensatory mechanisms. Nonetheless, the present results indicate that these variations of N-glycan maturation in AD may directly impact synaptic transmission.

N-glycan type and exact composition can have a significant impact on detection efficacy depending on conditions used for MS analysis, raising the concern that high levels of core-glycans observed in brain samples may be in part due to systematic experimental biases. High Man5 prevalence has, however, been reported in direct tissue - tissue comparisons with the same MS techniques (9) and has been confirmed with fluorescence HPCL, an assay that does not require glycan ionization and fragmentation (8). Although the actual proportion of core-glycans among neuronal proteins may thus vary from one MS setup to another, available data indicate that Man5, Man6, Man7 and Man8 constitute the largest proportion of N-glycans found in the total brain, the neuronal surface and synapses (Figure 1 and Supplemental Figure 1).

As depicted in textbooks, nascent glycoproteins typically exit the endoplasmic reticulum (ER) as Man8 forms (15). Man8 is then sequentially converted to Man7, 6 and Man5 in the medial/trans-Golgi where Man5 is modified by addition of N-acetylglucosamine (GlcNAc) on Man5 “A” branch by MGAT1, enabling further maturation into pauci, hybrid and complex N-glycan types in the medial or trans-Golgi (Figure 1A). According to this model, the most abundant N-glycans found at the neuronal surface and synapses thus appear to be processed no further than the medial/trans-Golgi. Underlying mechanisms are not clear at present. One possible explanation is that, due to their gigantic surface area, neuron needs for membrane protein synthesis exceed the upper bound of N-glycan processing capacity in the Golgi apparatus. As shown in mice and humans, MGAT1 mRNA appears to be downregulated in the brain compared to other tissues (25), which could potentially contribute to limit this capacity. However, this lower expression in the brain seems to be a general trend for glycosylation enzymes and is not specific to proteins controlling the maturation of core-glycans (9, 25).

Another possibility is that the subcellular organization of the neuronal N-glycosylation machinery may differ from what is typically observed in other cell types (26). Indeed, specific steps of core-glycan processing steps thought to primarily take place in the Golgi apparatus may also occur in other organelles, in particular the ER to Golgi intermediate compartments (IC). As directly shown in non-neuronal cell lines for ER endomannosidase and Golgi O-glycan N-acetylgalactosamine transferases (27, 28), glycosylation enzymes cycle between the ER or the Golgi and the IC and can accumulate in the later compartment under certain conditions (29). Our previous work and that of other groups on *endogenous proteins* of the secretory pathway have shown that dendrites, which constitute a large fraction of neuron volume, typically do not contain generic Golgi membranes but contain a substantial amount of ER and IC membranes (26). From the dendritic IC, nascent neuronal proteins can directly be sent to the neuronal surface *via* recycling endosomes, hence bypassing the Golgi apparatus located in neuron cell body (30). It will thus be interesting for future studies to determine whether Man8-6 can be directly processed into Man5 in the IC, which, given the large amounts of IC membranes in neurons, would provide a plausible cellular mechanism to account for the atypically high prevalence of Man5 at the neuronal surface.

We show here that core-glycans slow the desensitization of AMPA receptor complexes, thus demonstrating that distinct receptor glycotypes, rather than just the presence or absence of N-glycan at specific glycosylation sites, regulate synaptic signaling. More work will be needed to characterize underlying molecular mechanisms. One can envision that distinct N-glycans may have a direct and specific impact on structural changes occurring in core AMPA receptor subunits during desensitization. A significant challenge to addressing this hypothesis is that, owing to the very high conformational freedom of N-glycans (31), the structure of the extensively N-glycosylated domains of AMPA receptors complexes has not yet been elucidated. More tools are becoming available to model and elucidate the structure of whole glycoproteins (31), which will hopefully contribute to overcoming this challenge. Another and not mutually exclusive possibility is that distinct receptor glycotypes may regulate the association of core receptor subunits with distinct auxiliary subunits, in particular TARP γ8 (32–34) and Shisa6 (35), which are both known to regulate receptor desensitization and stabilization at synapses in hippocampal neurons. As indicated by recent MS, cryo-EM and electrophysiological studies, the composition of AMPA receptor complexes is much more diverse than initially thought (21, 36–38). Variability in receptor N-glycosylation likely contributes to and further augments this diversity. It will thus be important to determine whether specific glycotypes are associated with and favor the formation of specific compositions of AMPA receptor complexes.

Previous studies have shown that chronic (39) or acute (40) inhibition of N-glycan maturation in the Golgi impair LTP at CA3-CA1 synapses. However, these experiments were performed using organism-wide gene knockout or pharmacological inhibitors and it was insofar unclear whether LTP impairment was due to neuron specific processes. We show here that neuron specific inhibition of N-glycan maturation does not prevent the bulk of AMPA receptor trafficking at synapses but markedly reduce synaptic NMDA receptor signaling. More work will be required to determine whether this reflects an impairment of NMDA receptor expression at synapses and/or an alteration of their physiological properties. In this context, it is possible that core-glycans may have distinct and specific effects on AMPA and NMDA receptor complex assembly and itinerary through the neuronal secretory pathway (41). The present study did not address why neurons maintain high levels of surface N-glycan types that may limit a widespread form of postsynaptic plasticity and thereby, may impair learning. We have previously shown that core-glycosylated AMPA receptor glycotypes have a faster turnover than their standard glycoform counterparts (6). It will be interesting for further studies to determine whether this faster turnover and the slower desensitization of core-glycosylated AMPA receptor complexes that we describe here facilitate other and probably more labile forms of neuronal plasticity important for adaptative fitness.

## Material and methods

### Mgat1flx/flx mice

The mutant mice that were used in this study were generated by Marth and colleagues by insertion of *loxP* sites on both flanking sites of the single coding exon of the *Mgat1* gene (16). Heterozygous Mgat1^wt/flx^ mice (B6.129-*Mgat1^tm2Jxm/wt^*/J) were purchased from the Jackson Laboratory (JAX strain ID 006891). The transgenic line was maintained by inbreeding of homozygous animals in a hybrid B6/Swiss background. All animals were group housed in a standard 12hr light/12hr dark cycle. All experimental procedures were approved by the Animal Experimentation Ethical Comity of Université Paris Cité and authorized by the French Ministry of Higher Education, Research and Innovation.

### Drugs and AAVs

TTX, APV, gabazine (SR-95531) and NBQX were purchased from HeloBio. CGP56999A was purchased from Tocris. Sw was purchased from Toronto Research Chemicals-LGC Standards.

AAVs encoding GFP or mCherry± CRE recombinase scAAV-9/2-hSyn1-EGFP-SV40p(A), ID 256-9; ssAAV-9/2-hSyn1-mCherry-WPRE-hGHp(A), ID 253-9; ssAAV-9/2-hSyn1-chI-EGFP_2A_iCre-WPRE-SV40p(A), ID 146-9 and ssAAV-9/2-hSyn1-chI-mCherry_2A_iCre-WPRE-SV40p(A), ID 147-9) were purchased from the Viral Vector Facility of the University of Zurich.

### Primary neuronal cultures

Dissociated hippocampal neurons were prepared and maintained as described previously (6). In brief, primary cultures were prepared from P0-P1 wild-type Sprague-Dawley rat and Mgat1^flx/flx^ mouse pups after enzymatic and mechanical dissociation of dissected hippocampi. For imaging experiments, rat neurons were plated at a density of 13 to 19 x10^3^ cells/cm^2^ on glass coverslips coated with 333 µg/mL poly-D-lysine (PDL, Merck/Sigma-Aldrich #A-003-E). Mouse neurons were plated at a density of 13 to 19 x10^3^ cells/cm^2^ on glass coverslips coated with 333 µg/mL PDL and 100 to 200 µg/mL laminin (Merck/Sigma-Aldrich #L2020). For biochemical experiments, rat neurons were plated at a density of 26.2 x10^3^ cells/cm^2^ in plastic cultured dishes coated with 100 µg/mL PDL. Mouse neurons were plated at a density of 40x10^3^ cells/cm^2^ in culture dishes coated with 100 µg/mL PDL and 35 to 50 µg/mL lamimin. Neurons were maintained at 37°C in a 5% CO_2_ atmosphere in Neurobasal A medium (Gibco #10888022) supplemented with B27 (Gibco #11530536) and previously conditioned on confluent rat or mouse hippocampal glial cell cultures.

### Pharmacological and genetic inhibition of N-glycan maturation in cultured neurons

Rat hippocampal neuron cultures were treated with 30 µg/mL Swainsonine (Sw), a Golgi mannosidase2 inhibitor, for 7 days between DIV 13 and DIV 21.

Mgat1^flx/flx^ mouse hippocampus neuron cultures were transduced at DIV1 with adeno-associated virus (AAVs, see description above) encoding CRE and GFP or mCherry under a synapsin1 promoter, or as controls GFP or mCherry alone. Neurons plated on glass coverslips or in plastic dishes were transduced with 3.14 10^7^ viral genome (vg)/cm^2^, respectively, at DIV 1. Imaging, electrophysiological recordings and biochemical experiments were performed 3 to 4 weeks post-transduction.

### Stereotaxic injections of AAVs

Genetic inhibition of N-glycan maturation in CA1 hippocampal neurons *in vivo* was performed by unilateral stereotaxic injections of AAVs encoding GFP alone or GFP + CRE under a synapsin1 promoter (see AAV description below) in the dorsal hippocampus of 6 to 9 weeks-old Mgat1^flx/flx^ mice. Injections were performed at a flow rate of 100nL/min and the injection cannula was removed 5 minutes after infusion was complete. For spine morphology analysis, 300nL of PBS containing of 3.4 x10^11^ vg/mL were injected at the following stereotaxic coordinates (in mm from Bregma): AP: -1.9; ML, 1.3; at a depth (Z) of 1.5 (mm) from the brain surface. For electrophysiological recordings, 500 to 750nL of PBS containing 1.7 x10^12^ vg/mL were injected at (in mm): AP, -2.0; ML, 1.0; Z, 1.5.

### Antibodies, Nissl stain and lectins

The following antibodies were used for immunofluorescence labeling (IF) or immunoblotting (IB) at the indicated dilutions. Mouse anti-βactin (Sigma, IB, 1:10,000), rabbit anti-GFP (IF, Invitrogen, 1:2000), chicken anti-GFP (IF, Aves Labs, 1:5000), rabbit anti-GluA1 (IB, Abcam, 1: 2000; IF, Synaptic Systems, 1:1000), rabbit anti-GluA2 (IB, Abcam, 1:2000; IF, Synaptic systems, 1:1000)), rabbit anti-CRE (IF, Millipore, 1:1000), guinea-pig anti-VGLUT1 (IF, Sigma, 1:5000), mouse anti-MAP2 (IF, Sigma, 1:1000), IRDye secondary antibodies (IB, Li-Cor, 1:15,000), goat anti-guinea pig-Alexa647 (IF, Life Technologies, 1:750), goat anti-mouse (GAM)-Alexa594 (IF, Invitrogen, 1:500), goat anti-rabbit-RRX and GAM-FITC (Jackson Laboratory, ICC, 1:800). Nissl stain (NeuroTrace 640/660, Invitrogen, 1:2000) was used and added to secondary antibody solutions to label and identity neurons in some IF experiments in brain slices.

The following lectins (biotin conjugates) were used for far-western blotting at the indicated final concentrations: Concanavalin A (biotin-ConA, Sigma; 1 μg/mL), wheat germ agglutinin (biotin-WGA, Sigma; 1 μg/mL) and imaged by binding of IRDye-streptavidin (Licor, 1:15,000).

### Surface biotinylation, purification of surface proteins and deglycosylation

Surface biotinylation experiments were performed at room temperature essentially as described previously (6). In brief, 3 to 4 weeks-old neurons were rinsed in E4 medium (150 mM NaCl, 3 mM KCl, 15 mM glucose, 10 mM HEPES, [pH 7.4]) supplemented with 2 mM CaCl_2_, 2 mM MgCl_2_. Cells were then biotinylated for 10-12 min in E4 containing 1.25 mg/mL NHS-SS-Biotin (Pierce) and quenched in E4 containing 20mM glycine (Sigma-Aldrich). Cells were then scrapped in cold PBS supplemented with protease inhibitors (Protease inhibitor cocktail set III, EDTA-free, Merck/Sigma-Aldrich) supplemented with glycine and isolated by centrifugation. Cell pellets were then frozen for short-term storage or directly lysed and processed.

For standard immunoblotting experiments, cell pellets were lysed by trituration and incubation at 75°C (20min) in PBS containing 1% triton x100 and 0.5% SDS and supplemented with 20mM glycine, protease inhibitors and benzonase (Merck/Sigma-Aldrich). Biotinylated proteins were then purified as described previously (6) with sub-saturating amounts of high capacity streptavidine-agarose beads (Pierce) or streptavidine-magnetic particles (Pierce) in PBS supplemented with 1% Triton and 0.15% SDS and protease inhibitors, extensively washed and eluted in 0.5% SDS 1% Triton in PBS by reduction with 50-75mM DTT.

For mass spectrometry analyses, cell pellets were lysed by trituration and incubation at 75°C (20min) in PBS containing 2% SDS and supplemented with 20mM glycine, protease inhibitors and benzonase (Merck/Sigma-Aldrich). Biotinylated proteins were then purified as described previously (6) with sub-saturating amounts of high capacity streptavidine-agarose beads (Pierce) or streptavidine-magnetic particles (Pierce) in PBS supplemented with 0.25% SDS and protease inhibitors, extensively washed and eluted in 0.5% SDS in PBS by reduction with 50-75mM DTT.

For molecular shift assays, total or purified surface proteins were deglycosylated with peptide-N-glycosidase F (PNGase, New England Biolab) or endoglycosidase H (EndoHf, New England Biolab), which were used according to the manufacturer’s instructions. Proteins were typically digested for 3 hours at 37°C with 1000 (PNGase) or 3000 (EndoHf) units/ug protein.

### N-glycan profiling by MALDI-TOF-MS

MALDI-TOF-MS analyses of purified neuronal surface proteins were performed as described in ref. (42) with minor modifications.

#### N-glycan preparation

400ug of lyophilized protein lysate equivalent were resuspended in 0.6M TRIS buffer pH 8.5 and carbamidomethylated with iodoacetamide at 50°C for 1 hour. The samples were then dialyzed against 50 mM ammonium bicarbonate at 4°C for 16h-24h. N-glycans were then detached from proteins by overnight incubation with PNGase F at 37°C. The reaction was stopped by addition of a few drops of 5% acetic acid. Samples were washed on Sep-Pak C18 columns (50 mg, Waters) and collected in 5% acetic acid, dried in a speed-vac and lyophilized.

Lyophilized N-glycan samples were resuspended in 200 µL of a freshly prepared NaOH/DMSO slurry (7 NaOH pellets, Sigma-Aldrich #S8045; grounded in ∼3mL DMSO) and 100 µL of iodomethane (Sigma Aldrich, #289566). Samples were tightly capped and placed on a vortex shaker for 30 min at room temperature. After the mixture became white, semi-solid and chalky, 200 µL of ddH_2_O was added to stop the reaction and dissolve the sample. 200 µL of chloroform and an additional 400 µL ddH_2_O were added for chloroform extraction and vortexed followed by brief centrifugation. The aqueous phase was discarded and the chloroform fraction was washed three additional times with 800 µL ddH_2_O. Chloroform was then evaporated in a speed-vac. Permethylated glycans were resuspended in 200 µL of 50% methanol and added to a C18 Sep-Pak (50 mg) column preconditioned with methanol, ddH_2_O, acetonitrile, and ddH_2_O in succession. The reaction tubes were washed with 1 mL 15% acetonitrile and added to the column, followed by an additional 2 mL wash of 15% acetonitrile. Columns were placed into 15 mL glass round-top tubes, and permethylated glycans were eluted with 3 mL 50% acetonitrile. The eluted fraction was placed in a speed vacuum to remove the acetonitrile and lyophilized overnight.

#### MALDI-TOF-MS and N-glycan profiling

Permethylated glycans were resuspended in 25 µL of 75% methanol and spotted in a 1:1 ratio with DHB matrix on an MTP 384 polished steel target plate (Bruker Daltonics #8280781). MALDI-TOF MS data were acquired from a Bruker Ultraflex II instrument using FlexControl software in the reflective positive mode. A mass/charge (*m/z*) range of 1000–5000 kD was collected. Data was exported in MSD format for subsequent annotation. Glycans of known structure corresponding to the correct isotopic mass which had a signal to noise ratio greater than 6 (S/N) were annotated using mMass software (43) and their relative abundance was calculated as the signal intensity for each isotopic peak divided by the summed signal intensity for all measured glycans.

### N-glycan profiling by LC-ESI-MS/MS

#### N-glycan preparation

Surface protein samples were carbamidomethylated in 100 mM ammonium bicarbonate buffer and proteins precipitated with methanol/chloroform. Samples were redissolved in 30 µl 100 mM ammonium bicarbonate and N-glycan release by digestion with PNGase F at 37°C overnight. Released glycans were dried in a speed-vac and resuspended in 90 µL 100 mM ammonium bicarbonate buffer. 10 µL of a 500mM sodium borohydride solution was added and incubated overnight at room temperature.

N-glycans were purified using HyperSep Hypercarb SPE 10 mg cartridges (Thermo Scientific). The samples were applied to the cartridges, washed twice with an ammonium formate solution. Glycans were eluted with 500 µl of SPE elution solution, dried in a speed vac and redissolved in 20 µl ddH_2_O

#### LC-ESI-MS and N-glycan profiling

N-glycan samples were loaded on a porous graphitic carbon column (100 x 0.32 mm, Thermo Scientific) using 10 mM ammonium bicarbonate buffer as the aqueous solvent (solvent A) and acetonitrile as the eluant (solvent B). A steep gradient from 2 to 42 % of solvent B (98 to 58 % of solvent A) was developed over 17 min at a flow rate of 6 μl/min using an Ultimate 3000 capillary flow LC system (Thermo Scientific). Detection was performed with an ion trap mass spectrometer (Bruker amazon speed ETD) equipped with a standard ESI source in positive ion, DDA mode (= switching to MSMS mode for eluting peaks). MS-scans were recorded (range: 400-1600 Da). Data interpretation and quantification was performed with DataAnalysis 4.2 and QuantAnalysis 2.2 software (Bruker, Bremen, Germany). The extracted ion chromatogram of the first 4 isotopic mass peaks of every detected glycoform was integrated. Individual glycan species were quantified by quantifying corresponding total monoisotopic area (44).

### Retrospective analyses of published MS datasets

Analyses of published datasets were made with custom code in python 3 in a Jupyter Notebook environment (Anaconda3) using standard bioinformatics libraries (numpy, panda, seaborn, scipy, re, matplotlib).

#### N-glycan classification

Analyses were restricted to extended published MS datasets (see references in Main Text and Figure legends). For N-glycan classification, datasets were standardized and only information about N-glycan HexNAc, Hexoses, Fucoses, NeuAc (which were merged with NeuGC) and phosphates was kept. Glycans with 1 or less HexNAc were considered as degradation products; glycans with exactly 2 HexNAc and no phosphates, fucose nor NeuAc were classified as “ER” glycans if they had 8 or more Hex, and “ERGIC” if they had 5-7 Hex. Glycans with 3-4 Hex or other monosaccharide residues were classified as “HPC glycans”. Glycans with more than two HexNAc were classified as HPC glycans if they had at least 3 Hex, and were considered degraded otherwise.

#### Hierarchical clustering

Clustering heatmaps were generated with ClustVis 2.0 (45) with original data from ref. [7] with glycopeptides seen at least 20 times. N-glycan and protein information were clustered based on Euclidean distances using Ward’s method.

#### Gene ontology (GO)

GO analyses were performed using g:Profiler (46) by comparing protein clusters (*i.e.* “Man5” glycoproteins) using mouse annotated genes. Significance threshold was set to 0.01 and the g: scs threshold multiple test correction was applied.

### Immunofluorescence labeling

#### Cultured neurons

Cells were fixed in 4% PFA for 15 min at RT, blocked in PBS supplemented with 2% fish skin gelatin (FSG, Sigma) and 0.3% Triton X100 for 30 min and incubated overnight at 4**°**C with primary antibodies diluted in PBS supplemented with 1% FSG. Cells were then extensively washed in PBS, incubated with secondary antibodies diluted in PBS/FSG for 50 min, washed again and mounted in ProLong Diamond (Invitrogen). For labeling with rabbit anti-GluA1, cells were fixed by sequential 1min incubations in methanol and acetone previously cooled at -20**°**C in place of PFA.

#### Brain slices

Mouse brains were collected after intracardial perfusion of 4% PFA under deep anesthesia/analgesia, post-fixed by overnight incubation in 4% PFA at 4**°**C, washed in PBD and kept at 4**°**C in PBS/Azide until further processing. 50-70 μm coronal slices were prepared with a Leica VT1200S vibratome and kept at 4**°**C in PBS/azide until further processing. Slices were quenched with for 30 in 0.3 M glycine in PBS, rinsed in PBS and permeabilized with 0.2% Triton in PBS for 2-4 hours. Slices were then incubated in PBS supplemented with 3% normal goat serum (NGS, Invitrogen) and 0.2% Triton overnight at 4°C and then in primary antibodies diluted in the same buffer at 4°C till the next day. Slices were then washed in PBS supplemented with 0.2% triton, incubated with secondary antibodies diluted in PBS/NGS/triton overnight at 4°C, wash in PBS/triton and then PBS and mounted in ProlongDiamond.

### Confocal microscopy and image analysis

#### Imaging of cultured neurons

3D confocal imaging was performed using a x63 objective (NA=1.4) on a custom Leica/Yokogawa spinning disk setup (Errol) equipped with a sCMOS Flash4.0v3 Hamamatsu camera. Fluorescent pictures were analyzed using custom routines implemented in Metamorph (Molecular Devices) and Icy (47). In brief, maximal projections of picture z-series were denoised by low-pass filtering and segmented by wavelet decomposition (48). Synaptic AMPA receptors were quantified by measuring the integrated fluorescence intensity of receptor puncta with a 40-100% overlap with presynaptic elements identified by Vglut1 accumulation. Experimenters were blind to manipulations of N-glycosylation during recordings and analyses.

#### Imaging of brain slices and 3D reconstructions

3D confocal imaging was performed using x20 (NA=0.75) or x93 (NA=1.3) objectives on a Leica SP5 or a Leica SP8/STED3X microscope. Dendrites and dendritic spines 3D reconstruction and morphological analyses were performed with semi-automated routines using NeuroLucida360 (MBF Bioscience). Experimenters were blind to manipulations of N-glycosylation during recordings and analyses.

### Electrophysiological recordings in neuron cultures

Whole cell patch clamp recording was performed in 18-27 DIV neurons with an axopatch770B amplifier (Axon Instruments). For experiments in Mgat1^flx/flx^ mouse neurons, GFP fluorescence was used to identify AAV transduced neurons. The patch pipettes (3–4 MΩ) were pulled from borosilicate glass tubing and contained (in mM) 135 KMethylSulfate, 5 KCl, 0.1 EGTA-Na, 10 HEPES, 2 NaCl, 5 ATP, 0.4 GTP,10 phosphocreatine (pH 7.2; 280–290 mOsm). The extracellular solution contained (in mM) 125 NaCl, 2.5 KCl, 10 glucose, 26 NaHCO3, 1.25 NaH2PO4,2 Na Pyruvate, 2 CaCl2 and 1 MgCl2 and was saturated with 95% O2 and5% CO2, pH 7.3. Membrane potentials were corrected for liquid junction potential, which was measured to be ∼3 mV. Series resistance (typically less than 10 MΩ) was monitored and compensated throughout each experiment with the amplifier circuitry. 0.5 μM TTX and APV was bath-applied following dilution into the external solution from concentrated stock solutions. Outside out membrane patches were obtained by slow withdrawal of the recording electrode in whole cell patch clamp mode until a GΩ seal was re-formed. Patch pipettes were positioned upstream of a patch pipette with 1 MΩ resistance that was filled with 100 µM glutamate with equal concentration NaOH to maintain neutral pH and attached to a picoliter pressure injector (Warner Instuments). 10 ms pressure pulses were delivered to elicit AMPA currents in the outside-out patch pipette. During all experiments, the flow of the chamber was maintained at 3 mL/min. All data were acquired with Clampex v10.7 software (Molecular Devices) and analyzed with AxoGraph X (AxoGraph) and Igor pro (WaveMetrics). Experimenters were blind to manipulations of N-glycosylation during recordings and analyses.

### Electrophysiological recordings in acute brain slices

Transverse hippocampal slices were prepared 3 to 6 weeks after surgery. Animals were anesthetized with ketamine (20 mg/kg), xylazine (1.4 mg/kg) and isoflurane, and perfused transcardially with a modified sucrose cutting solution containing the following (in mM): sucrose 110, NaCl 10, KCl 2.5, NaH2PO4 1.25, NaHCO3 30, HEPES 20, glucose 25, thiourea 2, Na-ascorbate 5, Na-Pyruvate 3, CaCl2 0.5, MgCl2 10. Brains were then rapidly removed, hippocampi were dissected and placed upright into an agar mold and cut into 400 μm thick transverse slices in the same oxygenated cutting solution at 4°C using a Leica VT1200S vibratome. Slices were then immersed and kept in carbogen (95% O2/5% CO2)-bubbled artificial cerebrospinal fluid (ACSF) containing the following (in mM): NaCl 125, KCl 2.5, NaH2PO4 1.25, NaHCO3 26, glucose 10, Na-pyruvate 2, CaCl2 2, MgCl2 1. Slices were incubated at 32 °C for approximately 20 min then maintained at room temperature for at least 1 h. Prior to recording, slices were transferred to a recording chamber perfused with warm (32.5°C) oxygenated ACSF. Recording of field EPSPs (fEPSPs) was performed in current-clamp mode with a recording patch pipette (3–5 MΩ) containing 1 M of NaCl and positioned in the middle of *stratum radiatum* (SR) of hippocampal CA1 region with high densities of GFP expressing neurons. fEPSPs were evoked by mono-polar stimulation with a tip-broken-patch pipette filled with ACSF and positioned in CA1 SR. Pulses were delivered at 20-s intervals using Clampex, and voltage set using a Digitimer DS2A isolated stimulator to obtain 30-40% of fEPSP maximum amplitude. LTP was induced using a Theta-Burst Stimulation (TBS) protocol consisting of 4 stimulation trains delivered with 20-s intervals, with each train consisting of 5 bursts (5 pulses at 100 Hz per burst) interspaced by 200 ms. All fEPSPs recordings were performed in the presence of gabazine (1 μM, SR-95531) and CGP56999A (2 μM) to block GABA-A and GABA-B receptor mediated inhibitory transmission.

Data were recorded with a Multiclamp 700B amplifier (Axon Instruments) and acquired with Clampex. The slope of the field EPSPs was measured using Clampfit with all values normalized to a 10-min baseline period. The magnitude of plasticity was estimated by comparing averaged responses at 5–10 min after the induction protocol and 40–50 min after induction protocol. Results are reported as mean ± SEM.

For whole-cell recordings, neurons in the CA1 region were visually identified with infrared and GFP fluorescence imaging using an upright microscope equipped with a 40x objective. Patch electrodes (5– 6 MΩ) were filled with a low-chloride solution containing (in mM): 135 cesium methylsulfonate, 5 KCl, 0.1 EGTA-Na, 10 HEPES, 2 NaCl, 5 ATP, 0.4 GTP, and 10 phosphocreatine (pH adjusted to 7.25 with CsOH, 280-290 mOsm). To evoke EPSCs, a stimulating electrode was positioned in CA1 SR. All whole-cell recording experiments were performed in the presence of gabazine (1 μM) and CGP56999A (2 μM). AMPA to NMDA EPSC amplitude ratios, was calculated from the peak current measured at -70 mV (“AMPA”) over the current measured at + 50 mV 70 ms after the start of the stimulus artifact. Recordings of NMDA EPSCs were performed at -70mV in the presence of NBQX (50 μM) to block AMPA receptors. Series resistance (between 12 and 18 MΩ) was monitored throughout each experiment and cells showing more than 15% of change were excluded from analysis.

As mentioned above, AAVs encoding GFP and GFP+CRE were injected unilaterally in the dorsal hippocampus. Recordings were obtained in GFP expressing pyramidal neurons in slices transduced with GFP+CRE and, as controls, either in non-transduced neurons in contra-lateral slices or in GFP expressing neurons in slices transduced with GFP alone. These two control groups were balanced equally and combined for comparison with recordings from neurons transduced with GFP+CRE.

### Statistical analyses

Data are presented as means ± SEM unless otherwise indicated. Statistical analyses were performed in Prism (GraphPad). The number of measured values and independent experiments used for quantification are indicated in the text or in the figure legends. Mann Whitney non-parametric test was used to compare two means. Data normality was assessed with Augustino-Person’s or Kolmogorov Smirnov’s tests. When data passed normality test, one-way ANOVAs and post hoc Tukey or Sidak’s multicomparison tests were performed to determine whether significant differences existed among all or preselected pairs of means across multiple conditions (i.e. N>2). Otherwise, multiple comparisons were assessed with Kruskal-Wallis and Dunn’s multiple comparison tests.

## Acknowledgements

We thank Dorian Miremont, Camille Penet, Aude Lesmelle, Ada Bernaus, Anais Pereira, Kafi Md Abdullah Al, Idil Yuksekel, Sabrina Jaza and Ludivine Therreau for excellent technical assistance. We thank Diana Zala and Zsolt Lenkei for their input on the manuscript. We thank the staff of the animal facility of the IPNP for their support. MALDI-TOF-MS analyses were performed at the Glycomics core facility of the Beth Israel Deaconess Medical Center (Harvard Medical School, Boston, MA, USA) and coordinated by Christopher Ashwood, whom we thank for his support and expertise. LC-ESI-MS/MS analyses were performed at the BOKU mass spectrometry core facility of the University of Natural Resources and Life Sciences (Vienna, Austria) and coordinated by Clemens Grünwald-Gruber, whom we thank for his support and expertise. Cell and tissue imaging was carried out at the NeurImag Imaging core facility of the IPNP. We thank NeurImag staff for their support. We thank the Leducq Foundation for supporting the purchase of NeurImag Leica SP8 confocal/STED microscope and the Bettencourt Foundation for supporting the purchase of NeurImag custom spinning disc microscope. Work in the laboratory of CH is supported by the Agence Nationale de la Recherche (#ANR-16-CE16-0009-01 and #ANR-21-CE16-0021-01). Work in the laboratory of RP is supported by the ANR (#ANR-21-CE16-0021-01, ANR-18-CE370020-01, and ANR-18-CE16-0006) and the *Fondation pour la Recherche Médicale* (FRM Équipes EQU202003010457).

## Author contribution

C.-L.Z, C.M., M.T., A.V., R.P. and C.H. prepared samples, collected data and performed analyses.

R.P. and C.H. provided resources.

C.-L.Z, RP and C.H. designed the experiments and wrote the manuscript.

---

**Supplemental figure S1:**
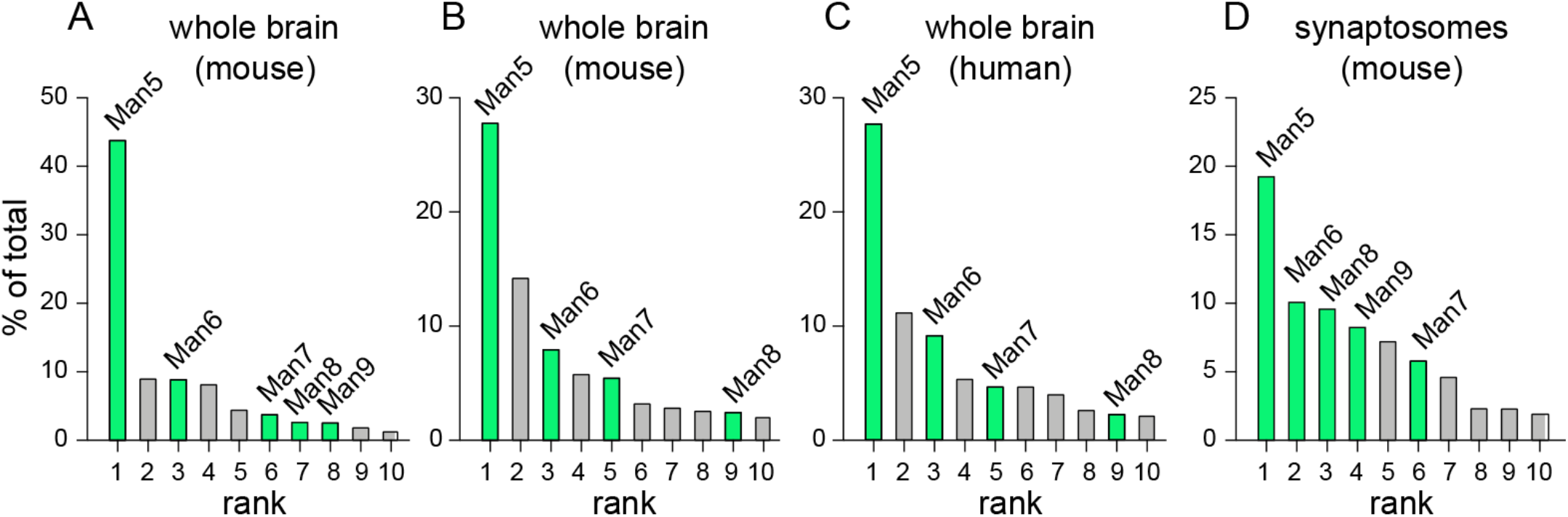
Prevalence of core-N-glycans in the brain and synapses, related to Figure 1. **A**-**D**) Occurrence of core-glycans (green) and HPC N-glycans (grey) among the top ten most abundant N-glycans identified by mass spectrometry (LC-MALDI-TOF MS) in total mouse brain (A and B, 2 independent studies), human brain (C) and mouse synaptosome samples (D). In the three types of samples, note the prevalence of Man5 and to a lesser extent Man6, Man8 and Man7. Plots were generated from data in ref. (9), (4), (4) and (8), respectively.

**Supplemental figure S2:**
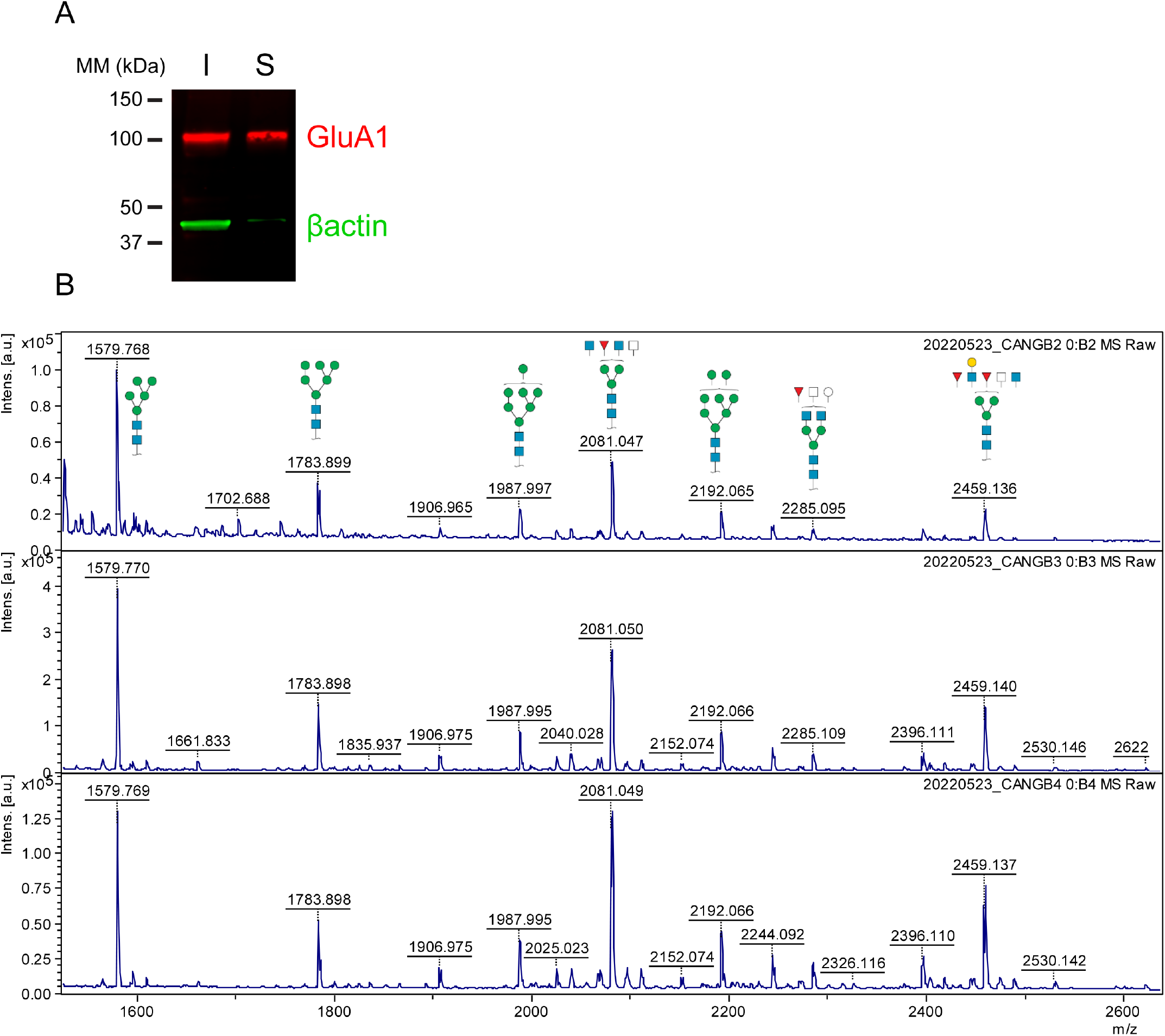
Prevalence of core-glycans at the neuronal surface, related to Figure 1. **A)** Immunoblot picture showing GluA1 (red) and βactin (green) in input and surface proteins (S) purified after surface biotinylation of cultured rat neurons. Note the exclusion of actin from surface samples and the presence of GluA1 in both input and surface fractions. **B)** m/z plots from 3 similar samples showing the most abundant N-glycans identified by LC-MALDI-TOF MS in neuronal surface proteins depicted in A.

**Supplemental figure S3:**
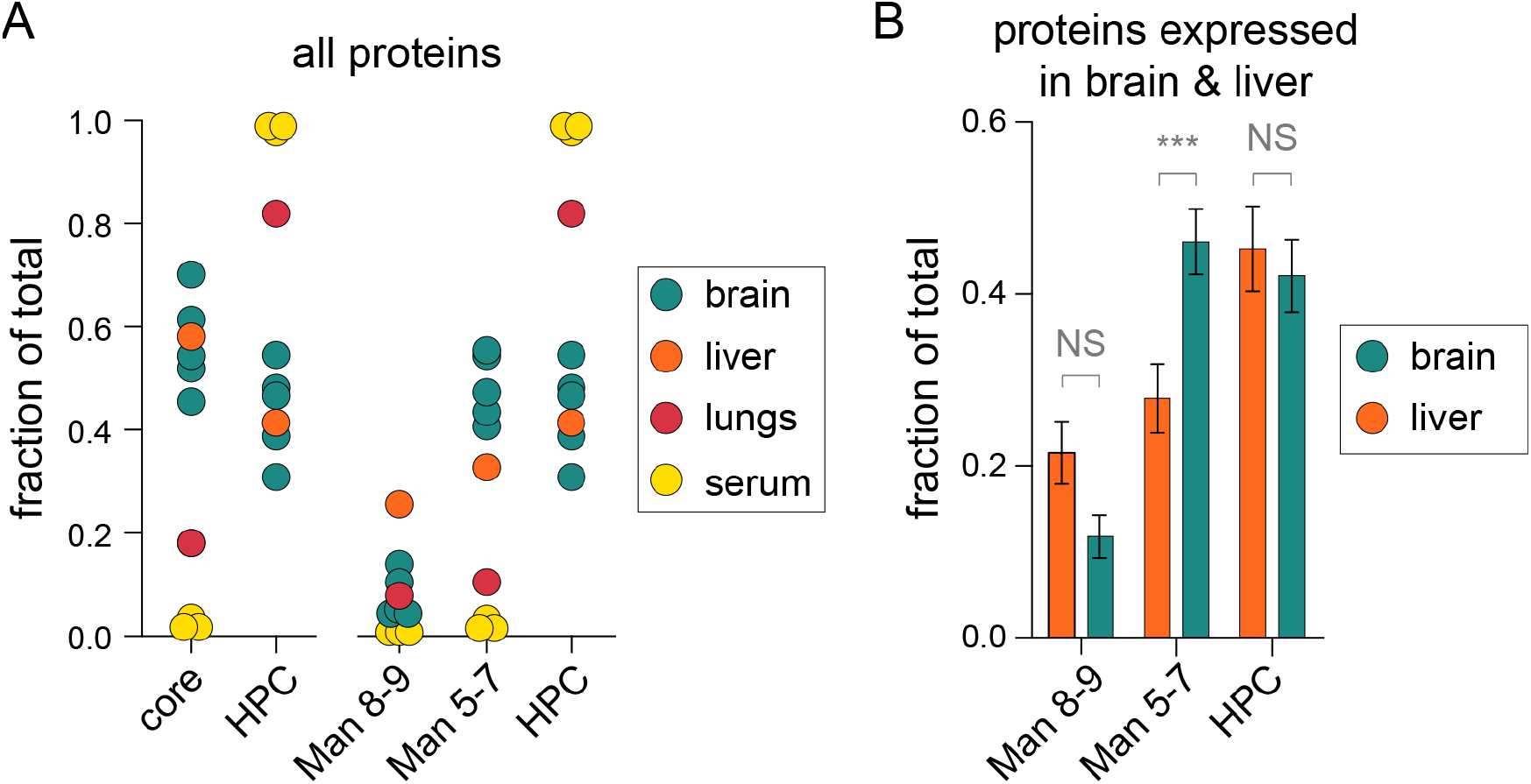
Prevalence of Man5-7 core-glycans is specific to the brain, related to Figure 1. **A)** Relative abundance of core and HPC N-glycans in mouse brain (5 studies), lung (1 study), liver (1 study) and serum (3 studies). Left-hand plot : comparison of total core-glycans *vs* HPC N-glycans showing that core-glycans are collectively particularly abundant in brain and in liver. Right-hand plot : comparison of Man8-9 *vs* Man5-7 *vs* HPC N-glycans showing that Man5-7, and not Man8-9, are more abundant in brain than liver. **B)** Relative abundance of Man8-9, Man5-7 and HPC N-glycans on proteins detected both in brain and liver (shown are means ± SEM, N=75), showing that the glycosylation of these proteins with Man5-7 is more frequent in the brain. Generated from raw data in ref. (4, 7–9, 13, 49–51). ***, p<10^-4^; Kruskal-Wallis’ and Dunn’s multi-comparisons tests.

**Supplemental Figure S4:**
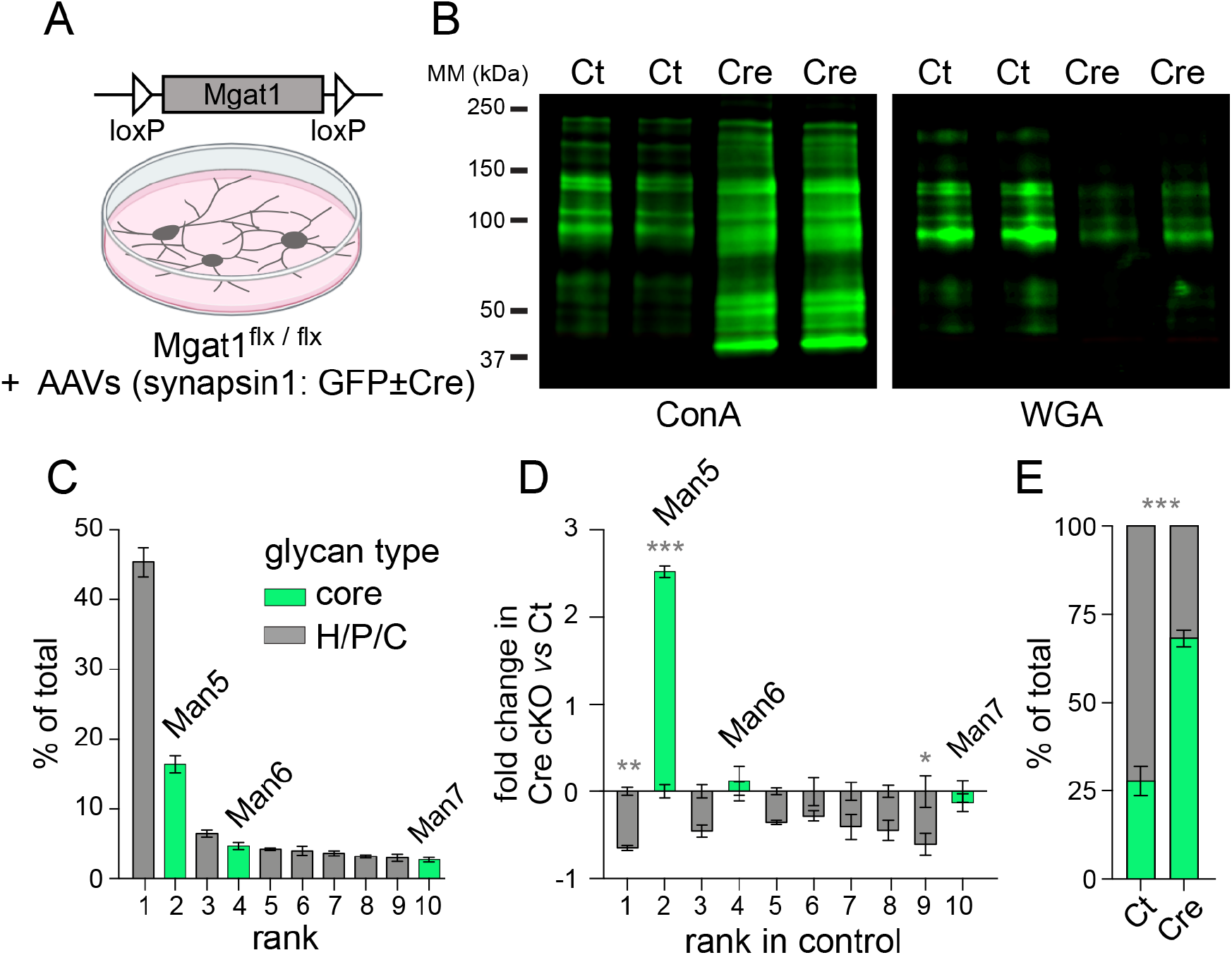
Conditional Mgat1 knock out enables experimental control of neuronal N-glycosylation, part 1, related to Figure 1. **A)** Transduction of cultured homozygous Mgat1^flx^ mouse neurons with Cre-encoding adeno-associated viruses (AAVs) enables conditional knockout (cKO) of Mgat1. **B**) Concanavalin A (ConA) and wheat germ agglutinin (WGA) labeling of far-western blots of surface proteins purified from cultured Mgat1^flx^ neurons transduced with GFP (control, Ct) or GFP+Cre AAVs (Cre) showing an increase of core-glycans (ConA labeling) and a decrease of complex glycans (WGA labeling) in Mgat1 KO samples. **C**) Occurrence of core-(green) and HPC N-glycan (grey) among the top 10 most abundant surface N-glycans (LC-ESI-MS) in control Mgat1^flx^ neurons. **D**-**E**) Relative expression of the same 10 N-glycans (D) and overall proportion of core-glycans (E) in Mgat1 KO versus control neurons (means ± SEM, n=3) showing a high expression of Man5 and increased levels of core-glycans at the expense of HPC N-glycans in Mgat1 KO neurons.

**Supplemental Figure S5:**
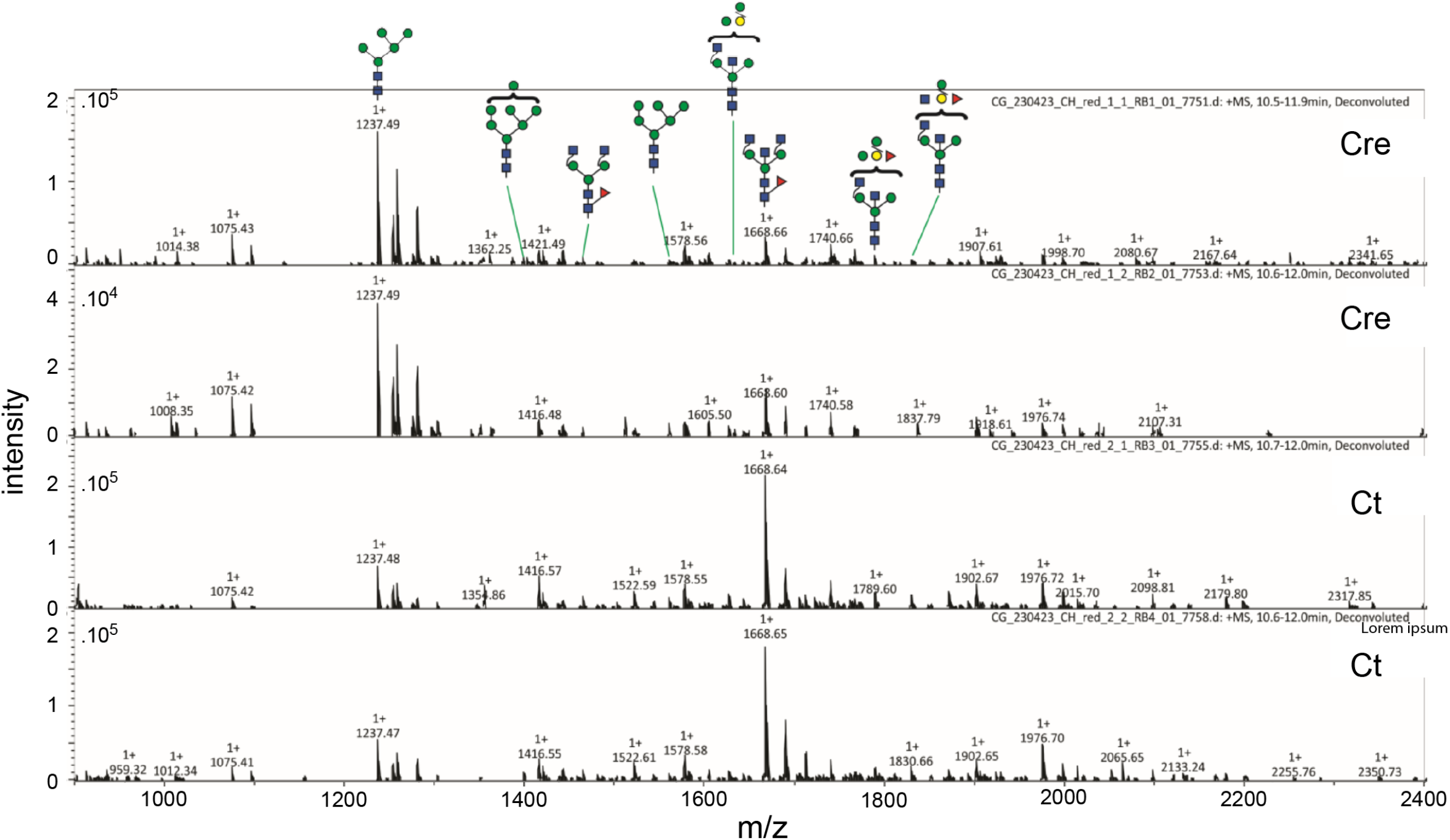
Conditional Mgat1 knock out enables experimental control of neuronal N-glycosylation, part 2, related to Figure 1. Examples of LC-ESI-MS spectra in control and Mgat1 cKO cultured neurons (see Supplemental Figure S4C-S4D).

**Supplemental Figure S6.**
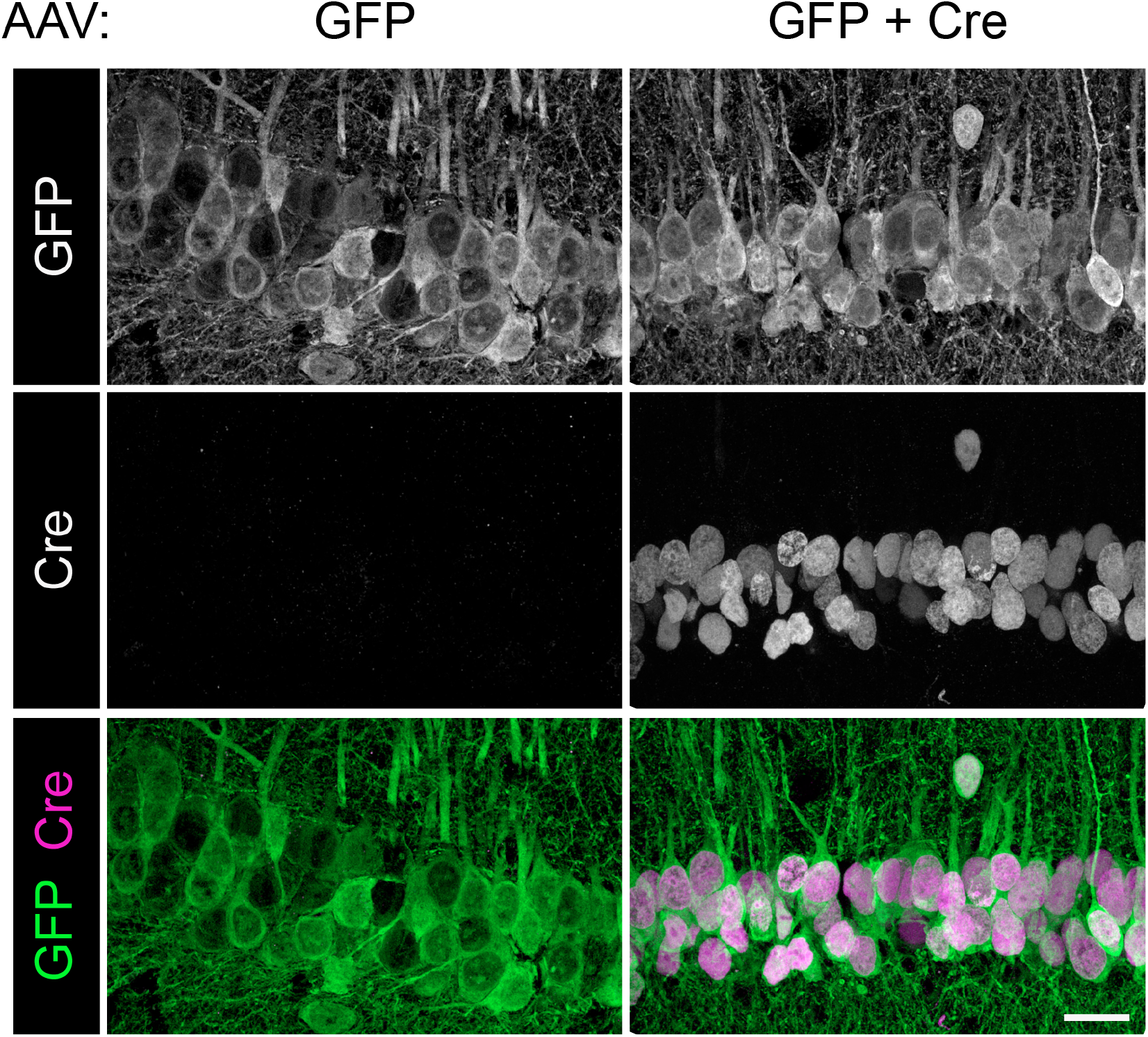
Cre expression after viral transduction of Mgat1^flx^ neurons *in vivo*, related to Figure 2. Confocal micrographs of Mgat1^flx^ hippocampal slices prepared and immuno-labeled for GFP and Cre 8 weeks after stereotaxic injection of AAV encoding GFP or GFP+Cre. Note the expression and nuclear localization of Cre in neurons transduced with “GFP+Cre” AAVa. Scale bar 25 microns.

**Supplemental Figure S7.**
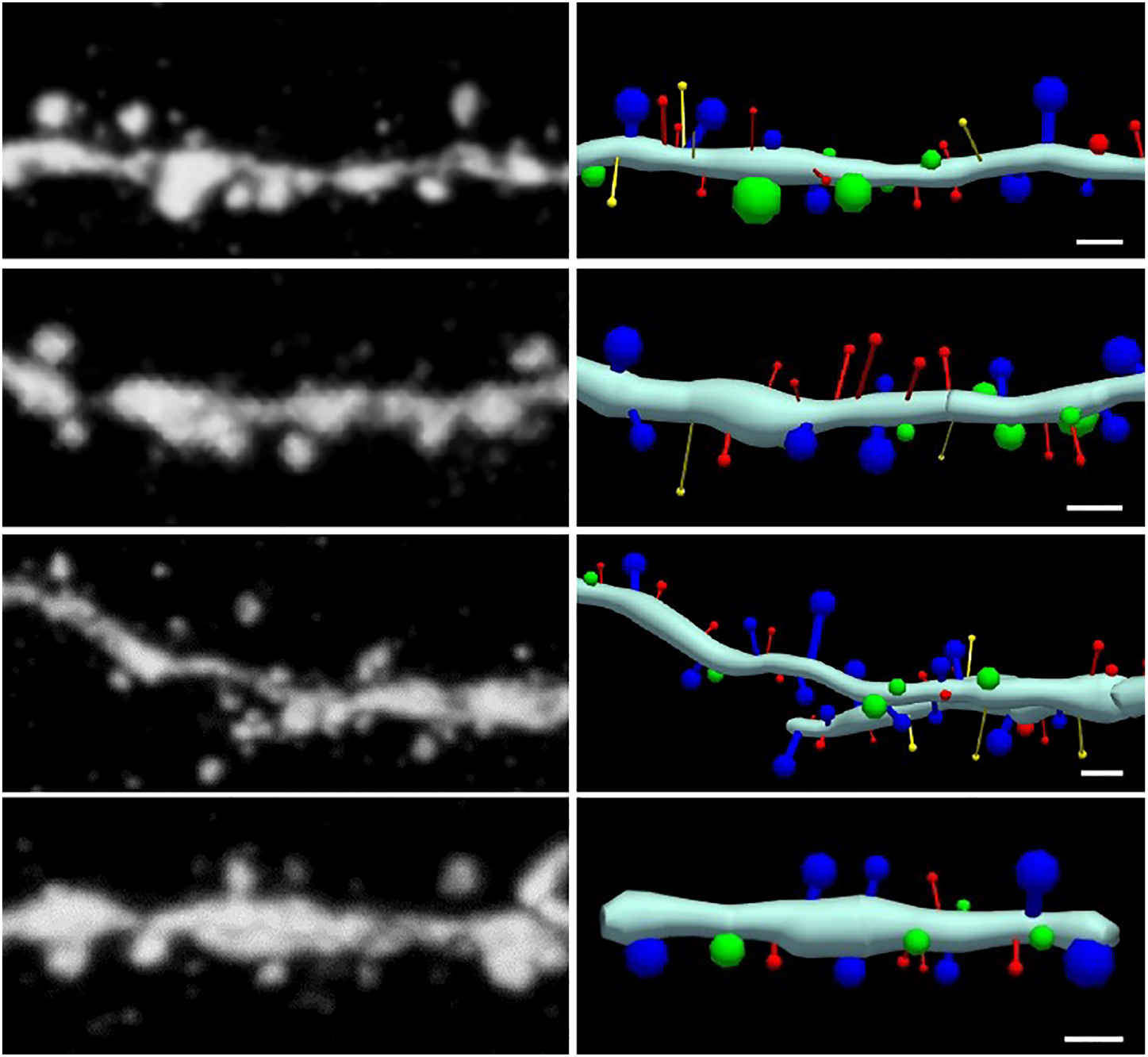
3D reconstruction of dendritic spines, related to Figure 2. Confocal micrographs and 3D reconstruction of GFP expressing pyramidal neuron secondary and tertiary dendritic branches in CA1 *stratrum radiatum*. Filopodia, thin, mushroom and stubby dendritic spines are shown in yellow, red, blue and green, respectively. Scale bar 5 microns.

**Supplemental Figure S8.**
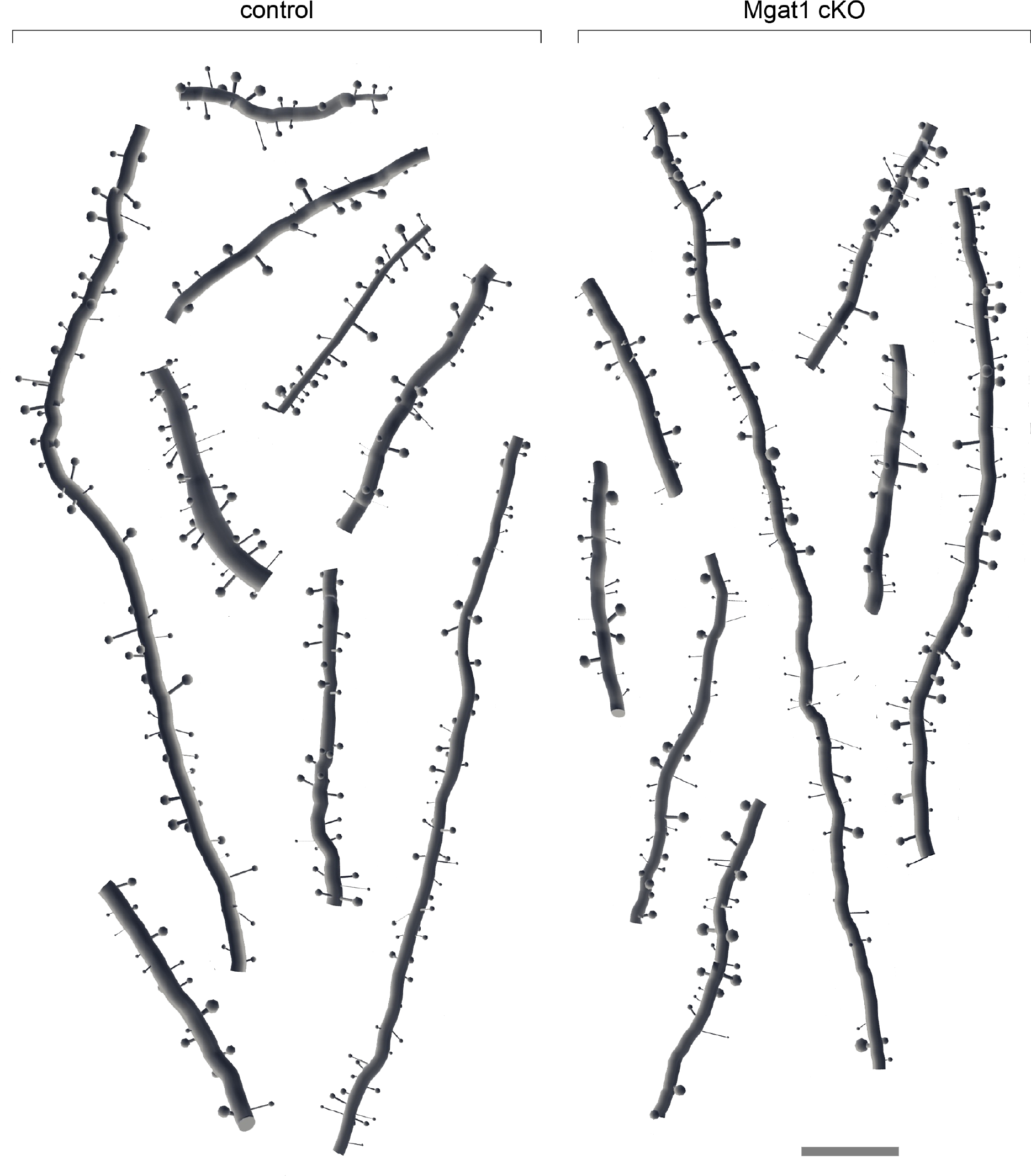
Example reconstructions of control (Ct) and Mgat1 cKO dendrites, related to Figure 2. Scale bar 5 microns.

**Supplemental Figure S9.**
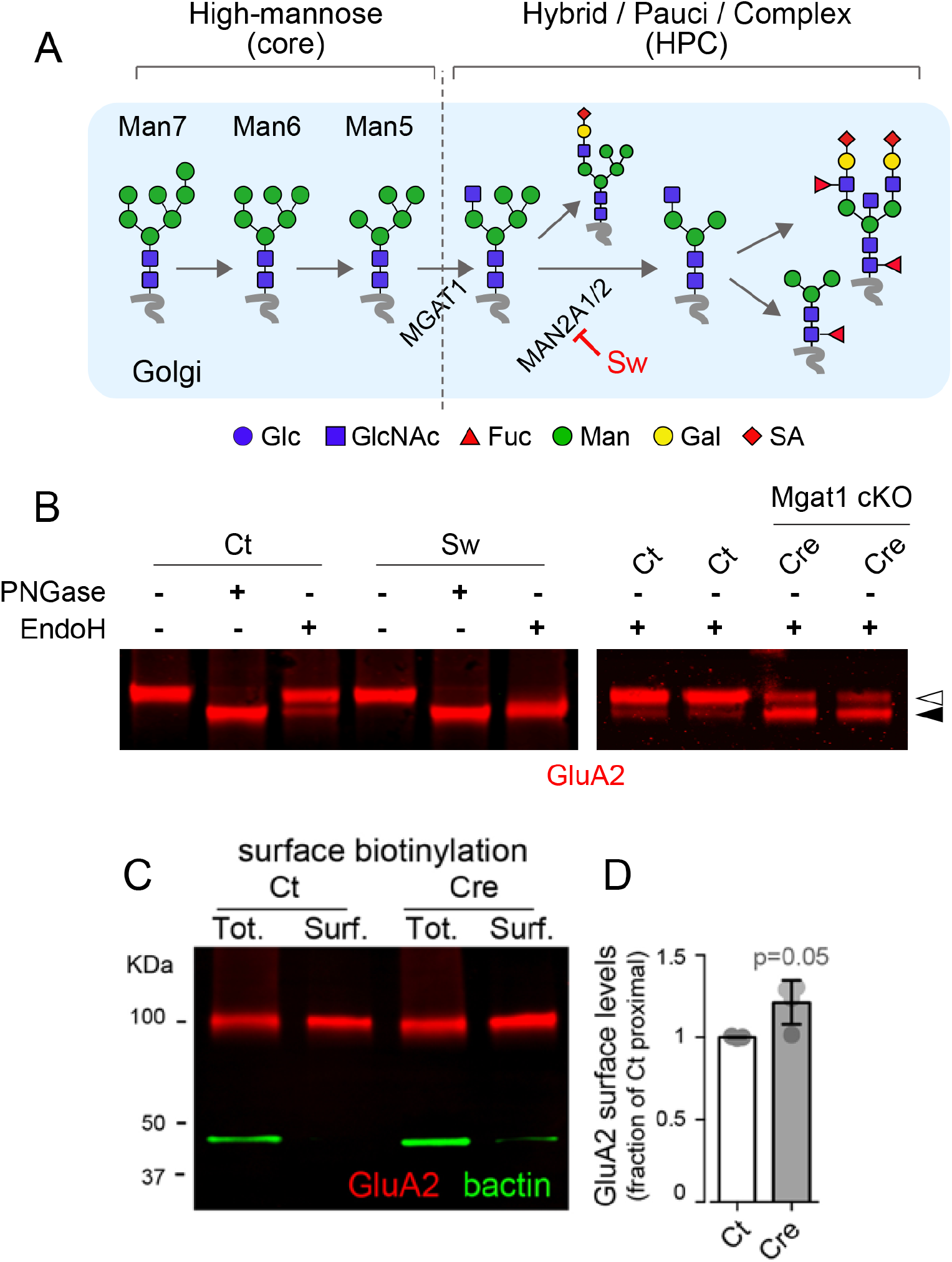
Core-glycosylation does not impair AMPA receptor expression at the neuronal surface, related to Figure 2. **A)** Simplified scheme of N-glycan processing in the Golgi apparatus highlighting steps blocked by Swainsonine (Sw) and MGAT1 cKO. Glc, glucose; GlcNAc, N-acetyl-glucosamine; Fuc, fucose; Man, mannose; Gal, galactose; SA, sialic acid. **B)** Immunoblot pictures of surface AMPA receptors (GluA2, red) from control neurons (Ct), neurons chronically treated with swainsonine (Sw) and Mgat1 cKO neurons (Cre), before or after deglycosylation with PNGase (removal of all N-glycans) or EndoH (removal of core-glycans only). In both pictures, note the increased levels of EndoH sensitive GluA2 after inhibition of N-glycan maturation (filled arrowhead). **C,D**) Immunoblot pictures (B) and quantification (C) of GluA2 in total cultured neuron lysates (Tot.) and purified surface protein fractions (Surf.) in control conditions (Ct) or Mgat1 KO (Cre) showing that inhibition of core-glycan maturation does not reduce AMPA receptor expression at the neuronal surface. Shown in B are means ± SEM. N=4 (p=0.05, *Mann-Whitney test*).

**Supplemental figure S10.**
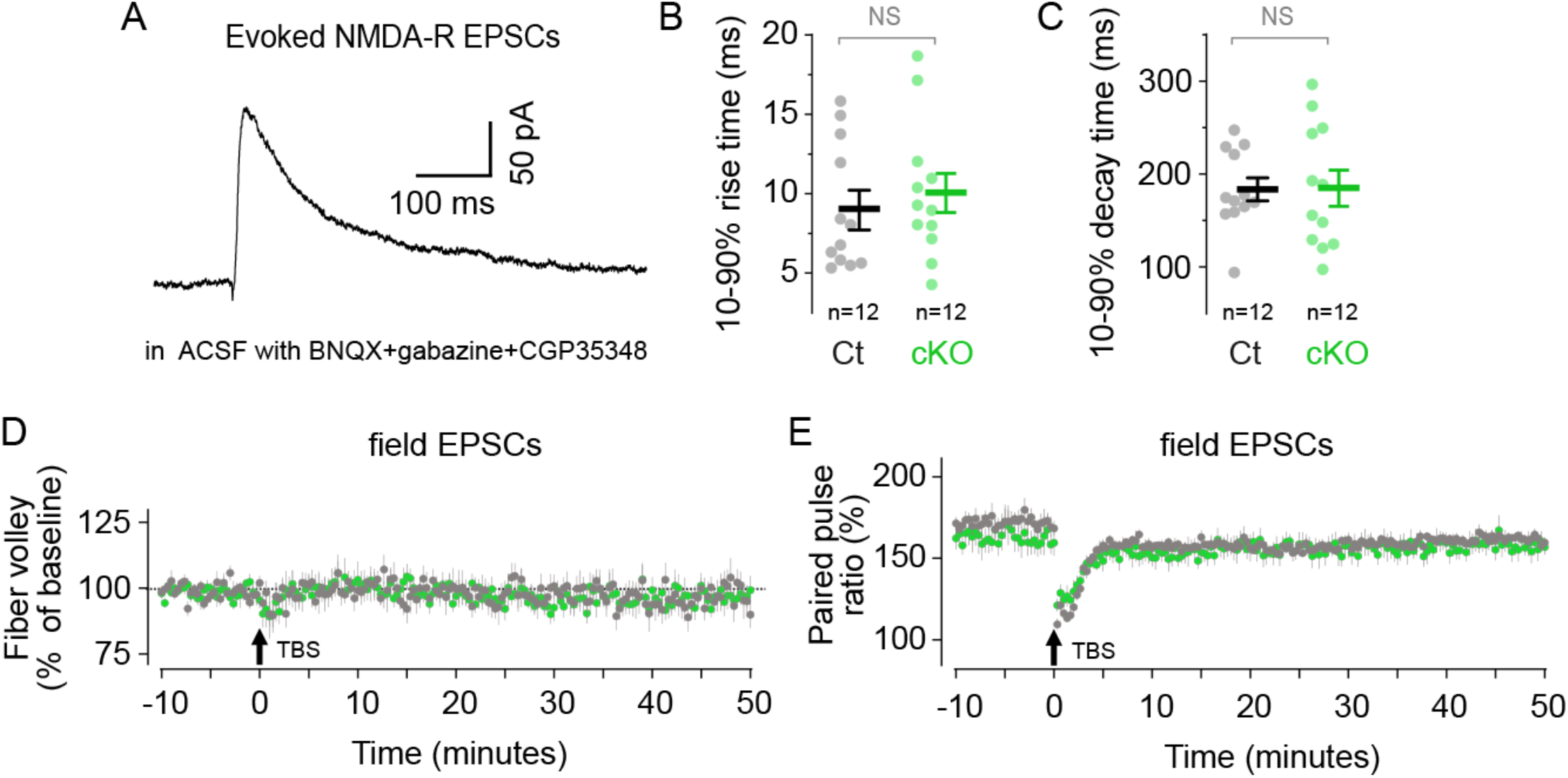
related to Figure 4. **A)** Example trace of evoked NMDA-R mediated EPSC. The top 10% to 90% rise time (B) and decay time (C) are not affected by cKO of Mgat1. N=12 recordings from 5-6 animals per group. Shown are means ± SEM. NS, not significant; Mann-Whitney’s test. **D**-**E**) Summary plots (means ± SEM) of normalized fiber volley amplitude and paired-pulse ratio (PPR) during fEPSC recording. N=7-9 slices from 5-8 animals per group.

## References

1. K. W. Moremen, M. Tiemeyer, A. V. Nairn, Vertebrate protein glycosylation: diversity, synthesis and function. Nat Rev Mol Cell Bio 13, 448–462 (2012).

2. T. Štambuk, M. Klasić, V. Zoldoš, G. Lauc, N-glycans as functional effectors of genetic and epigenetic disease risk. Mol. Asp. Med. 79, 100891 (2021).

3. S. E. Williams, R. G. Mealer, E. M. Scolnick, J. W. Smoller, R. D. Cummings, Aberrant glycosylation in schizophrenia: a review of 25 years of post-mortem brain studies. Mol Psychiatr 25, 3198–3207 (2020).

4. S. Suttapitugsakul, et al., Glycoproteomics Landscape of Asymptomatic and Symptomatic Human Alzheimer’s Disease Brain. Mol Cell Proteomics 21, 100433 (2022).

5. Q. Zhang, C. Ma, L.-S. Chin, L. Li, Integrative glycoproteomics reveals protein N-glycosylation aberrations and glycoproteomic network alterations in Alzheimer’s disease. Sci Adv 6, eabc5802 (2020).

6. C. Hanus, et al., Unconventional secretory processing diversifies neuronal ion channel properties. Elife 5, e20609 (2016).

7. N. M. Riley, A. S. Hebert, M. S. Westphall, J. J. Coon, Capturing site-specific heterogeneity with large-scale N-glycoproteome analysis. Nat Commun 10, 1311 (2019).

8. M. M. Bradberry, T. M. Peters-Clarke, E. Shishkova, E. R. Chapman, J. J. Coon, N-glycoproteomics of brain synapses and synaptic vesicles. Cell Reports 42, 112368 (2023).

9. S. E. Williams, et al., Mammalian brain glycoproteins exhibit diminished glycan complexity compared to other tissues. Nat Commun 13, 275 (2022).

10. A. R. Yale, et al., Regulation of neural stem cell differentiation and brain development by MGAT5-mediated N-glycosylation. Stem Cell Rep (2023). 10.1016/j.stemcr.2023.04.007.

11. A. R. Yale, et al., Cell Surface N-Glycans Influence Electrophysiological Properties and Fate Potential of Neural Stem Cells. Stem Cell Rep. 11, 869–882 (2018).

12. M. B. Kandel, et al., N-glycosylation of the AMPA-type glutamate receptor regulates cell surface expression and tetramer formation affecting channel function. J. Neurochem. 147, 730–747 (2018).

13. J. C. Trinidad, R. Schoepfer, A. L. Burlingame, K. F. Medzihradszky, N-and O-Glycosylation in the Murine Synaptosome*. Mol. Cell. Proteom. 12, 3474–3488 (2013).

14. J. R. Cooney, J. L. Hurlburt, D. K. Selig, K. M. Harris, J. C. Fiala, Endosomal Compartments Serve Multiple Hippocampal Dendritic Spines from a Widespread Rather Than a Local Store of Recycling Membrane. J. Neurosci. 22, 2215–2224 (2002).

15. A. Varki, et al., Essentials of Glycobiology, 4th edition (2022).

16. Z. Ye, J. D. Marth, N-glycan branching requirement in neuronal and postnatal viability. Glycobiology 14, 547–558 (2004).

17. K. Harris, J. Stevens, Dendritic spines of CA 1 pyramidal cells in the rat hippocampus: serial electron microscopy with reference to their biophysical characteristics. J. Neurosci. 9, 2982–2997 (1989).

18. M. Matsuzaki, N. Honkura, G. C. R. Ellis-Davies, H. Kasai, Structural basis of long-term potentiation in single dendritic spines. Nature 429, 761–766 (2004).

19. J. N. Bourne, K. M. Harris, Balancing Structure and Function at Hippocampal Dendritic Spines. Annu. Rev. Neurosci. 31, 47–67 (2008).

20. Y. Fukazawa, R. Shigemoto, Intra-synapse-type and inter-synapse-type relationships between synaptic size and AMPAR expression. Curr. Opin. Neurobiol. 22, 446–452 (2012).

21. J. Schwenk, et al., Regional Diversity and Developmental Dynamics of the AMPA-Receptor Proteome in the Mammalian Brain. Neuron 84, 41–54 (2014).

22. J. E. Huettner, Glutamate receptor channels in rat DRG neurons: Activation by kainate and quisqualate and blockade of desensitization by con A. Neuron 5, 255–266 (1990).

23. K. M. Partin, D. K. Patneau, C. A. Winters, M. L. Mayer, A. Buonanno, Selective modulation of desensitization at AMPA versus kainate receptors by cyclothiazide and concanavalin A. Neuron 11, 1069–1082 (1993).

24. G. L. Collingridge, S. J. Kehl, H. McLennan, Excitatory amino acids in synaptic transmission in the Schaffer collateral-commissural pathway of the rat hippocampus. J. Physiol. 334, 33–46 (1983).

25. X. Tang, et al., Transcriptomic and glycomic analyses highlight pathway-specific glycosylation alterations unique to Alzheimer’s disease. Sci. Rep. 13, 7816 (2023).

26. M. J. Kennedy, C. Hanus, Architecture and Dynamics of the Neuronal Secretory Network. Annu Rev Cell Dev Bi 35, 1–24 (2019).

27. C. Zuber, M. J. Spiro, B. Guhl, R. G. Spiro, J. Roth, Golgi Apparatus Immunolocalization of Endomannosidase Suggests Post-Endoplasmic Reticulum Glucose Trimming: Implications for Quality Control. Mol. Biol. Cell 11, 4227–4240 (2000).

28. T. Jarvela, A. D. Linstedt, Irradiation-induced protein inactivation reveals Golgi enzyme cycling to cell periphery. J. Cell Sci. 125, 973–980 (2012).

29. D. J. Gill, J. Chia, J. Senewiratne, F. Bard, Regulation of O-glycosylation through Golgi-to-ER relocation of initiation enzymes. J. Cell Biol. 189, 843–858 (2010).

30. A. B. Bowen, A. M. Bourke, B. G. Hiester, C. Hanus, M. J. Kennedy, Golgi-independent secretory trafficking through recycling endosomes in neuronal dendrites and spines. Elife 6, e27362 (2017).

31. Y.-X. Tsai, et al., Rapid simulation of glycoprotein structures by grafting and steric exclusion of glycan conformer libraries. Cell 187, 1296–1311.e26 (2024).

32. S. Chen, et al., Activation and Desensitization Mechanism of AMPA Receptor-TARP Complex by Cryo-EM. Cell 170, 1234–1246.e14 (2017).

33. B. Herguedas, et al., Mechanisms underlying TARP modulation of the GluA1/2-γ8 AMPA receptor. Nat. Commun. 13, 734 (2022).

34. N. Rouach, et al., TARP γ-8 controls hippocampal AMPA receptor number, distribution and synaptic plasticity. Nat. Neurosci. 8, 1525–1533 (2005).

35. R. V. Klaassen, et al., Shisa6 traps AMPA receptors at postsynaptic sites and prevents their desensitization during synaptic activity. Nat. Commun. 7, 10682 (2016).

36. Y. Zhao, S. Chen, A. C. Swensen, W.-J. Qian, E. Gouaux, Architecture and subunit arrangement of native AMPA receptors elucidated by cryo-EM. Science 364, 355–362 (2019).

37. N. P. Pampaloni, I. Riva, A. L. Carbone, A. J. R. Plested, Slow AMPA receptors in hippocampal principal cells. Cell Rep. 36, 109496 (2021).

38. J. Schwenk, et al., An ER Assembly Line of AMPA-Receptors Controls Excitatory Neurotransmission and Its Plasticity. Neuron 104, 680–692.e9 (2019).

39. W. Gu, et al., Loss of α1,6-Fucosyltransferase Decreases Hippocampal Long Term Potentiation implications for core fucosylation in the regulation of AMPA receptor heteromerization and cellular signaling. J. Biol. Chem. 290, 17566–17575 (2015).

40. H. Matthies, et al., Glycosylation of proteins during a critical time window is necessary for the maintenance of long-term potentiation in the hippocampal CA1 region. Neuroscience 91, 175–183 (1999).

41. O. Jeyifous, et al., SAP97 and CASK mediate sorting of NMDA receptors through a previously unknown secretory pathway. Nat. Neurosci. 12, 1011–1019 (2009).

42. R. G. Mealer, et al., The schizophrenia risk locus in SLC39A8 alters brain metal transport and plasma glycosylation. Sci. Rep. 10, 13162 (2020).

43. M. Strohalm, D. Kavan, P. Novák, M. Volný, V. Havlíček, mMass 3: A Cross-Platform Software Environment for Precise Analysis of Mass Spectrometric Data. Anal. Chem. 82, 4648–4651 (2010).

44. C. Grünwald-Gruber, A. Thader, D. Maresch, T. Dalik, F. Altmann, Determination of true ratios of different N-glycan structures in electrospray ionization mass spectrometry. Anal. Bioanal. Chem. 409, 2519–2530 (2017).

45. T. Metsalu, J. Vilo, ClustVis: a web tool for visualizing clustering of multivariate data using Principal Component Analysis and heatmap. Nucleic Acids Res. 43, W566–W570 (2015).

46. U. Raudvere, et al., g:Profiler: a web server for functional enrichment analysis and conversions of gene lists (2019 update). Nucleic Acids Res. 47, W191–W198 (2019).

47. F. de Chaumont, et al., Icy: an open bioimage informatics platform for extended reproducible research. Nat. Methods 9, 690–696 (2012).

48. C. Hanus, et al., Synaptic Control of Secretory Trafficking in Dendrites. Cell Reports 7, 1771–1778 (2014).

49. Y. Yoshida, et al., Quantitative analysis of total serum glycome in human and mouse. PROTEOMICS 16, 2747–2758 (2016).

50. N. Jia, et al., Glycomic Characterization of Respiratory Tract Tissues of Ferrets Implications for its Use in Influenza Virus Infection Studies*. J. Biol. Chem. 289, 28489–28504 (2014).

51. K. F. Medzihradszky, K. Kaasik, R. J. Chalkley, Tissue-Specific Glycosylation at the Glycopeptide Level* [S]. Mol. Cell. Proteom. 14, 2103–2110 (2015).

